# An antisense oligomer conjugate with unpredicted bactericidal activity against *Fusobacterium nucleatum*

**DOI:** 10.1101/2025.02.12.637808

**Authors:** Valentina Cosi, Jakob Jung, Linda Popella, Falk Ponath, Chandradhish Ghosh, Lars Barquist, Jörg Vogel

## Abstract

Fusobacteria are commensal members of the oral microbiome that can spread from their primary niche and colonize distal sites in the human body. Their enrichment in colorectal and breast cancer tissue has been associated with tumor growth, metastasis, and chemotherapeutic resistance. The use of non-selective antibiotics to remove fusobacteria impairs tumor progression, but prolonged application risks side effects such as gastrointestinal problems and dysbiosis. Species-specific antisense antibiotics based on peptide nucleic acid (PNA) have shown efficacy in many Gram-negative species, suggesting that antisense PNAs may also enable a tailored depletion of fusobacteria. Here, we have investigated the antibacterial potential of cell-penetrating peptide (CPP)-PNA conjugates targeting the mRNA of putative essential genes in *Fusobacterium nucleatum*. Unexpectedly, we observed no growth inhibition with any of the target-specific PNAs, but identified a non-targeting control CPP-PNA (FUS79, (RXR)_4_XB-GACATAATTGT) as a potent growth inhibitor of *F. nucleatum*. Our data suggest that the CPP and specific sequence features of FUS79 are responsible for its activity, rather than an antisense effect. Interestingly, FUS79 also inhibits the growth of five additional fusobacterial strains but not of *F. nucleatum* ssp. *vincentii* (FNV). RNA-seq analysis indicates that FUS79 induces a membrane stress response in a vulnerable *F. nucleatum* strain but not in FNV. Collectively, our attempt at developing antisense antibiotics for fusobacteria discovers a potent growth inhibitor, whose bactericidal effect appears independent of target-specific mRNA inhibition.

**IMPORTANCE:** Enrichment of *F. nucleatum* at cancer sites is associated with increased tumor growth and metastasis. Antibiotic treatment to remove the bacteria was shown to change the course of cancer progression. Here we explore first steps to establish peptide nucleic acids (PNAs) as specific antisense antibiotics, thereby laying the foundation for further development of antisense technology in fusobacteria. While the CPP-PNA FUS79 was initially designed as a control, we observed that the compound was bactericidal for specific fusobacterial strains. Our results suggest that FUS79 might be able to selectively deplete fusobacterial strains from bacterial communities, offering a new perspective on fusobacterial removal at the tumor site.

## INTRODUCTION

The human oral microbiome is composed of more than 700 bacterial species from seven different phyla, including the phylum Fusobacteriota (1). Fusobacteria are Gram-negative non-motile anaerobic bacteria found in the oral cavity and throat (2). They use their elongated shape and numerous adhesins to connect early and late colonizers of the oral niche (3–5), effectively functioning as a central bridging organism (6–8). One member of this phylum, *Fusobacterium nucleatum*, has received attention for its implication in medical conditions such as oral infections, arthritis, adverse pregnancy outcomes, and most importantly cancer (9). *F. nucleatum* can disseminate throughout the body via the gastro-intestinal or hematogenous route and colonize secondary sites (10, 11). Enrichment of *F. nucleatum* at tumor sites is associated with increased cell proliferation, metastasis, inhibition of immune responses and resistance to chemotherapy (12–14). Antibiotic treatment has been shown to counteract *F. nucleatum*-associated tumor growth (15, 16), but sustained administration of broad spectrum antibiotics runs the risk of severe side effects such as a systemic inflammatory response and disease reoccurrence (17). This motivates the development of *Fusobacterium*-specific antimicrobial agents for a selective removal of this oncomicrobe.

Antisense oligomers (ASOs) have great potential for the development of programmable species-specific antibiotics (18–21). These ‘asobiotics’ are typically designed to base pair with the translation initiation region (TIR) of mRNAs encoding an essential gene in the species of interest. Through antisense sequestration of the TIR, ASOs prevent synthesis of the targeted protein, resulting in bacterial growth inhibition. Two ASO modalities have been popular for asobiotics design: peptide nucleic acid (PNA) and the phosphorodiamidate morpholino oligomer (PMO). Both are neutral in charge and resistant to cellular nucleases and proteases (22). PNA, the more widely used of the two, is a synthetic DNA analog composed of a peptide-like backbone and natural nucleobases (23). Antisense PNAs composed of 9 to 12 nucleobases possess strong binding affinity for DNA and RNA (24) and have been shown to effectively inhibit translation of target transcripts in *Escherichia coli or Salmonella enterica* (25, 24). However, neither PNA nor PMO can passively cross the bacterial envelope (26, 27) and require a carrier for cell entry. The most common approach to facilitate PNA uptake is conjugation to cell-penetrating peptides (CPPs) (28), which are relatively easy to synthetize or modify (29). CPPs have been shown to deliver PNAs into various Gram-negative (30) as well as Gram-positive (31) bacteria. PNAs conjugated to the polymyxin-inspired peptide (KFF)_3_K or the arginine-rich peptide (RXR)_4_XB have demonstrated potent antimicrobial activity against the Gram-negative pathogen *S. enterica* with a typical minimal inhibitory concentration (MIC) in the lower micromolar range (32). PNAs have been extensively used against aerobic Gram-negative bacteria (33, 34). In addition, there is one report of their use against *Porphyromonas gingivalis* (35), an obligate anaerobic Gram-negative bacterium, with which *F. nucleatum* interacts in the oral cavity. However, PNA-mediated growth inhibition of fusobacteria remained to be tested.

Here, we have attempted to use PNAs as programmable antisense antibiotics to kill fusobacteria. We establish that the CPP (RXR)_4_XB readily enters *F. nucleatum*. Yet, we found that ASOs conjugated to (RXR)_4_XB targeting mRNAs of essential genes do not inhibit the growth of fusobacteria in culture. Surprisingly, a non-targeting (RXR)_4_XB-PNA conjugate (FUS79) displays potent antimicrobial activity against five different fusobacterial strains. This effect seems to be caused by the combination of the (RXR)_4_XB peptide and certain sequence elements of the conjugated ASO, rather than antisense inhibition of an off-target mRNA. Global RNA-seq analysis reveals that FUS79 elicits a membrane stress response and activates the σE regulon in sensitive fusobacteria but not in a resistant fusobacterial species. In summary, our results represent a first step toward the design of ASO therapeutics against *F. nucleatum* and tell a cautionary tale arguing that appropriate controls are needed in the development of CPP-ASOs as antimicrobial agents.

## RESULTS

### Cell-penetrating peptides (KFF)_3_K and (RXR)_4_XB enter F. nucleatum

To explore CPPs as potential PNA delivery agents in *F. nucleatum*, we coupled the fluorophore TAMRA to four different CPPs to investigate CPP-mediated uptake into the bacterial cytoplasm, a method previously applied in *E. coli* (36). Using the clinical isolate *F. nucleatum* ssp. *nucleatum* ATCC 23726 (FNN23) we determined uptake efficiency of the CPPs (KFF)_3_K, (RXR)_4_XB, Seq471 and Seq2373 coupled to TAMRA (Table S1). (KFF)_3_K and (RXR)_4_XB are known to deliver charge-neutral ASOs in numerous Gram-negative bacteria (19). Seq471 (WLRRIKAWLRRIKALNRQLGVAA) has a high transport efficiency into HeLa cells without notable cytotoxicity, making it an interesting candidate to target intracellular *F. nucleatum* (37). Seq2373 (AHKLKKPKIVRLIKFLLKAWK) was designed in-house and showed good preliminary uptake efficiency for FNN23 in a flow cytometry screen. Since many CPPs possess an inherent antibacterial activity against Gram-negative bacteria (38, 39), we first determined the MIC (minimum inhibitory concentration) of the CPP-TAMRA conjugates. Seq471 and Seq2373 displayed an MIC of 10 µM, whereas (KFF)_3_K and (RXR)_4_XB showed an MIC of 20 µM and >40 µM, respectively (Fig. S1A). We therefore tested CPP-TAMRA uptake at 5 µM to avoid toxicity.

Using confocal laser scanning microscopy (CLSM) we observed that the uptake of CPPs in FNN23 varies greatly (Fig. 1A). (RXR)_4_XB has strong and stable penetration efficiency with 100% TAMRA-positive cells from 10 minutes onwards. (KFF)_3_K showed ∼15% uptake efficiency with a slight decrease over time. Seq471 and Seq2373 showed minimal to no translocation with 0-4% TAMRA-positive cells (Fig. 1B). To confirm that our lead CPP (RXR)_4_XB delivers the fluorophore into the bacteria, we used z-stack recordings to detect (RXR)_4_XB-coupled TAMRA in the cytosol of FNN23 and not in the membrane, the latter judged by membrane staining with FM 4-64 (Fig. 1C).

**Fig. 1:**
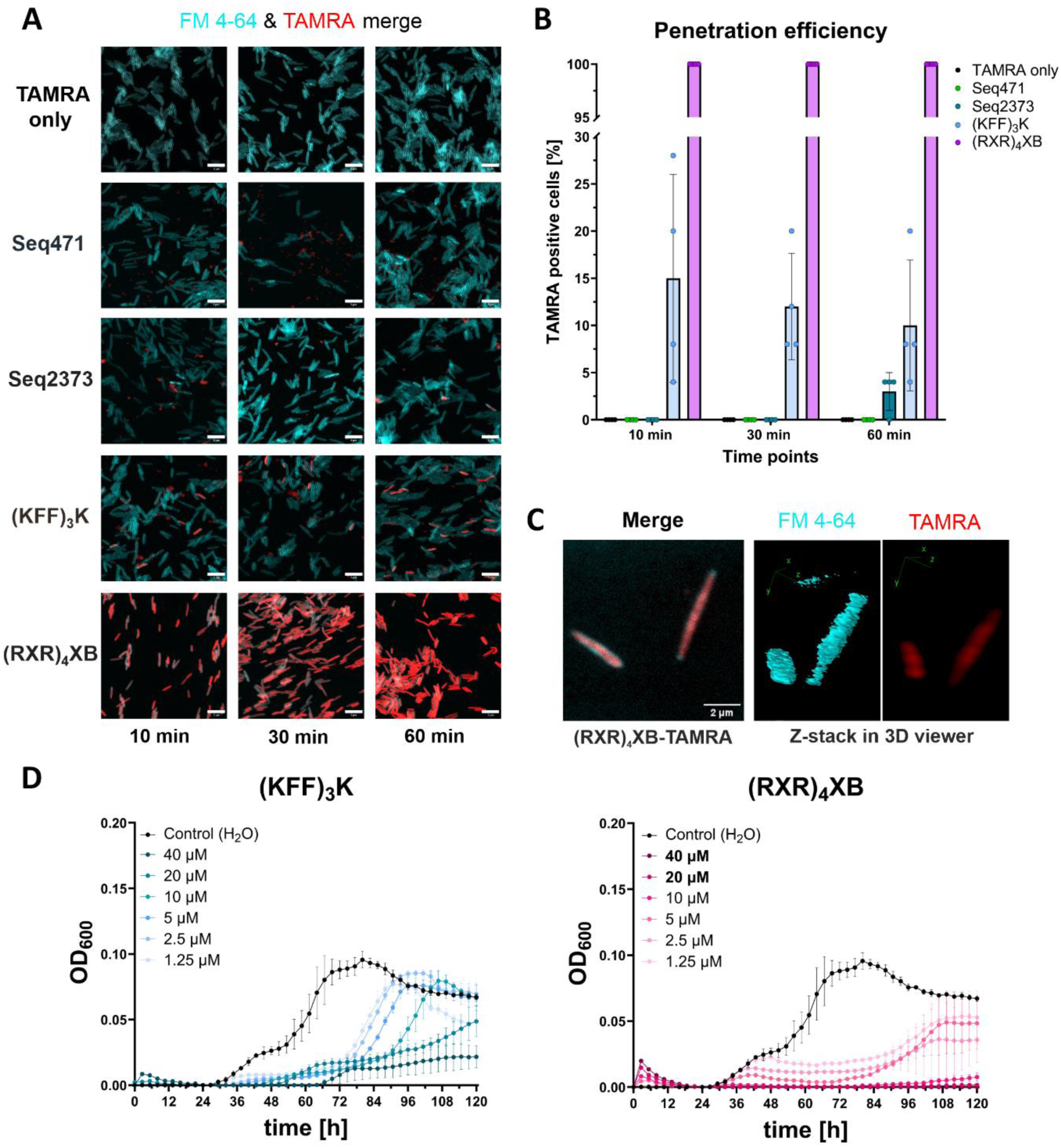
CPP (RXR)_4_XB is a potent delivery tool for *F. nucleatum*. (A) Representative CLSM images for TAMRA only control and TAMRA-labeled CPPs at 10, 30, and 60 minutes post treatment. FNN23 was incubated with FM 4-64 to stain the cell membrane, which is shown in cyan. Scale bar, 5 µm. (B) Quantification of TAMRA-positive cells for each CPP at selected time points. A total of 100 cells from four different images per condition were selected via bright field and cytoplasmic TAMRA signals were counted. Each dot represents the percentage of TAMRA positive cells from one image. Mean of four images is shown. Error bars indicate standard deviation of three experiments. (C) Z-stack of (RXR)_4_XB-TAMRA treated FNN23. IMAGE-J 3D viewer was used to represent z-stacks. (D) MIC of unconjugated CPPs using FNN23 in MHB at 1 x 10^5^ CFU per ml input. Growth curves are depicted as OD_600nm_ over time for three replicates from different o.n. cultures, error bars indicate standard deviation of three experiments.

Since TAMRA conjugation can alter the cytotoxicity of CPPs (40), we determined the MIC of unconjugated (KFF)_3_K and (RXR)_4_XB. The MIC of (KFF)_3_K was >40 µM (Fig. 1D), whereas (RXR)_4_XB showed an MIC of 20 µM (Fig. 1D). Since both CPPs exhibited antibacterial activity at higher concentrations, we selected 10 µM as maximal initial concentration for the following experiments.

### Design of CPP-PNA conjugates to inhibit protein synthesis of fusobacterial essential gene

To identify potential mRNA targets for inhibiting *F. nucleatum* growth, we initially selected three genes assumed to be essential based on inference from other bacterial species (25, 31, 41, 42): *acpP* (C4N14_04130), which encodes acyl carrier protein; *ftsZ* (C4N14_10675), which encodes the Z-ring protein that is important for cell division; and *gyrA* (C4N14_02325), which encodes the gyrase A subunit important for DNA replication (Fig. 2A). For each of these three genes, we designed 10mer PNAs complementary to the TIR using the previously published FNN23 annotation and the web-tool MASON (43–45). To account for unspecific effects, each PNA sequence was scrambled and tested together with the respective targeting PNA as a control.

**Fig. 2:**
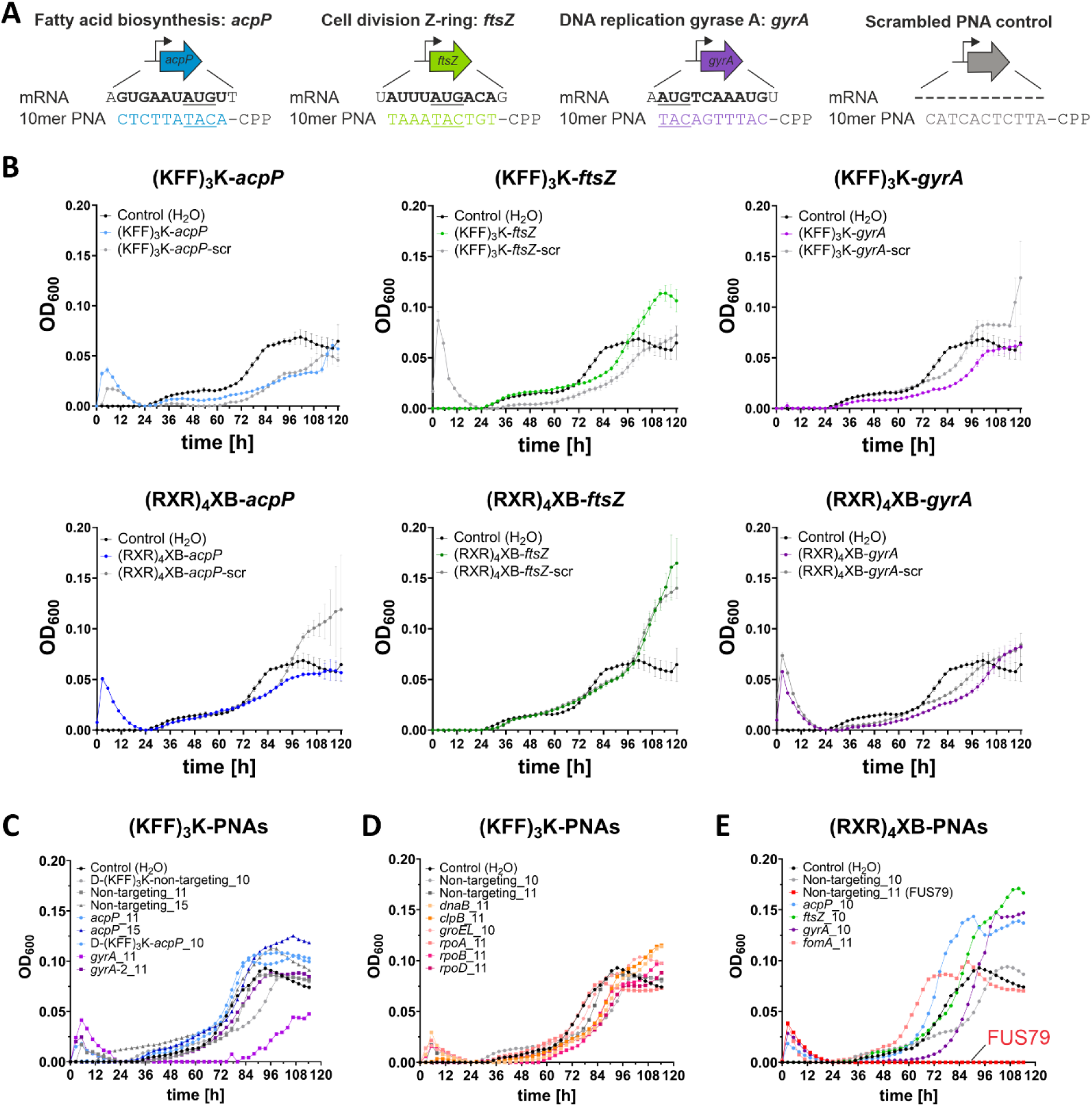
FNN23 growth is not inhibited by mRNA targeting CPP-PNAs, but by a non-targeting (RXR)_4_XB-PNA control. (A) Overview of PNA design for three gene targets in FNN23. 10mer PNAs were designed with sequence complementarity along the translational start codon region (‘AUG’ underlined) of the target mRNAs depicted in bold for *acpP* (blue), *ftsZ* (green) and *gyrA* (purple). For each PNA the sequence was scrambled as control. An example of a respective scrambled control is shown for the *acpP* sequence (gray). (B) Growth inhibition assays of FNN23 using 10 µM CPP-PNAs. Upper row shows (KFF)_3_K-PNAs. On-target PNAs are complementary to *acpP* (light blue), *ftsZ* (light green) and *gyrA* (lilac). Lower row shows (RXR)_4_XB-PNAs. On-target PNAs are complementary to *acpP* (dark blue), *ftsZ* (dark green) and *gyrA* (purple). Corresponding scrambled controls are depicted in gray. All growth curves are depicted as OD_600nm_ over time for three replicates from different o.n. cultures, error bars indicate the standard deviation of three experiments. (C) Growth inhibition of FNN23 with (KFF)_3_K-PNA alternatives for *acpP*, *ftsZ* and *gyrA*. Non-targeting (KFF)_3_K-PNA controls are shown in gray. D-form (KFF)_3_K CPPs are indicated respectively. (D) Growth inhibition assays of FNN23 using (KFF)_3_K-PNAs. Each color represents a different gene. Non-targeting (KFF)_3_K-PNA controls shown in gray. (E) Growth inhibition of FNN23 with (RXR)_4_XB-PNAs targeting alternative sites of *acpP*, *ftsZ* and *gyrA* or (RXR)_4_XB-PNA targeting *fomA*. Each color represents a different gene. Non-targeting (RXR)_4_XB-PNA 10mer control is shown in gray. Non-targeting (RXR)_4_XB-PNA 11mer control (FUS79) is shown in red. All growth curves were performed in MHB using 1 x 10^5^ CFU per ml and 10 µM CPP-PNAs. Growth is depicted as OD_600nm_ reading over time.

### Treatment of F. nucleatum with CPP-PNAs delays but does not block growth

To test CPP-PNA-conjugates for growth inhibition, we diluted mid-exponential phase cultures of FNN23 to 1 x 10^5^ colony forming units (CFUs) per ml in Mueller-Hinton Broth (MHB) and incubated the diluted bacteria with 10 µM of either (KFF)_3_K- or (RXR)_4_XB-conjugated PNAs targeting *acpP*, *ftsZ* or *gyrA*. Growth was monitored over time by OD_600nm_ measurements with growth inhibition defined as no visible growth (OD_600nm_ <0.01) after 120 hours. Unexpectedly, we did not observe growth inhibition from any CPP-PNA (Fig. 2B). Although some PNAs lead to growth delay, the effect did not seem to be due to target-specific translation inhibition because the respective scrambled controls showed comparable effects (Fig. 2B). Since *ftsZ* inhibition via CRISPRi was reported to prevent FNN23 colony formation on agar plates, but not in liquid culture (46), we tested if the (RXR)_4_XB-PNA targeting *ftsZ* had an effect on CFU numbers. However, we could not observe a CFU reduction after CPP-PNA treatment (Fig. S1B). Based on these findings we conclude that CPP-PNAs targeting fusobacterial mRNAs are not effective in antisense inhibition.

Next, we explored additional CPP-PNA variations (Table S2) based on (KFF)_3_K-PNAs, since the unconjugated CPP was less toxic to FNN23 compared to (RXR)_4_XB (Fig. 1D). Since PNA length affects efficacy (25, 47), we also tested 11mer and 15mer PNAs. In addition, we chose a different 10mer PNA sequence for *gyrA*, shifting the binding site downstream to test if this target window would be more efficient (-1 to +10 relative to AUG start codon; referred to as *gyrA*_2). In addition, we included a D-isomeric form of the lysine in the (KFF)_3_K peptide to avoid putative proteolytic cleavage (30, 48, 49). We used a single non-targeting (KFF)_3_K-PNA control for each PNA with similar length and nucleobase composition. These controls had no predicted binding to any TIR of an annotated FNN23 gene. Of these (KFF)_3_K-PNAs, only the 11mer *gyrA*-2 PNA caused a substantial growth delay of ∼40 hours (Fig. 2C). Higher concentrations of this CPP-PNA did not lead to further growth retardation and CFU spot assays showed a similar effect compared to the non-targeting control (Fig. S1C and S1D). This suggests that the observed inhibition is most likely mediated by an unspecific effect caused by the CPP-PNA conjugate and not by *gyrA* inhibition.

We then selected eight additional target genes and designed (KFF)_3_K-conjugated 10mer and 11mer PNAs (Table S2) targeting mRNAs encoding proteins involved in DNA transcription (*dnaB*, *rpoA*, *rpoB* and *rpoD*) or stress chaperones (*clpB* and *groEL*), because some of these targets inhibited growth in uropathogenic *E. coli* (25), *Listeria monocytogenes* (50) and *P. gingivalis* (35). None of these eight additional PNAs inhibited fusobacterial growth (Fig. 2D).

### Screening of additional (RXR)_4_XB-PNAs leads to the discovery of FUS79

Since (RXR)_4_XB showed better uptake efficiency in the initial TAMRA screen compared to (KFF)_3_K, we also tested (RXR)_4_XB-conjugated PNAs targeting alternative sequences. Specifically, we designed PNAs targeting sequences around the TIR of *acpP*, *ftsZ* or *gyrA* as well as an 11mer PNA targeting *fomA,* a very abundant outer membrane protein of fusobacteria (43, 51). As controls, we included non-targeting 10mer and 11mer (RXR)_4_XB-PNAs to account for unspecific effects. None of the on-target PNAs impaired growth (Fig. 2E). However, the non-targeting (RXR)_4_XB-PNA 11mer control (i.e., (RXR)_4_XB-GACATAATTGT, named FUS79 from here on) completely inhibited growth of FNN23 (Fig. 2E, shown in red). This was unexpected as this ASO sequence was specifically designed to lack binding sites within TIRs of FNN23 and no other (RXR)_4_XB-PNA conjugate had shown growth inhibition at similar concentrations.

### FUS79 is bactericidal against F. nucleatum

To determine the MIC of FUS79, we performed a growth inhibition assay with a dilution series of this PNA. FUS79 leads to complete growth inhibition of FNN23 with an MIC of 10 µM, whereas a scrambled FUS79 sequence (referred from here on as PNAscr) had no impact on growth (Fig. 3A). To exclude a potential batch effect, we reordered FUS79 from the original vendor and also initiated an in-house synthesis. Both batches confirmed the FUS79-mediated growth inhibition (Fig. S2A and S2B).

**Fig. 3:**
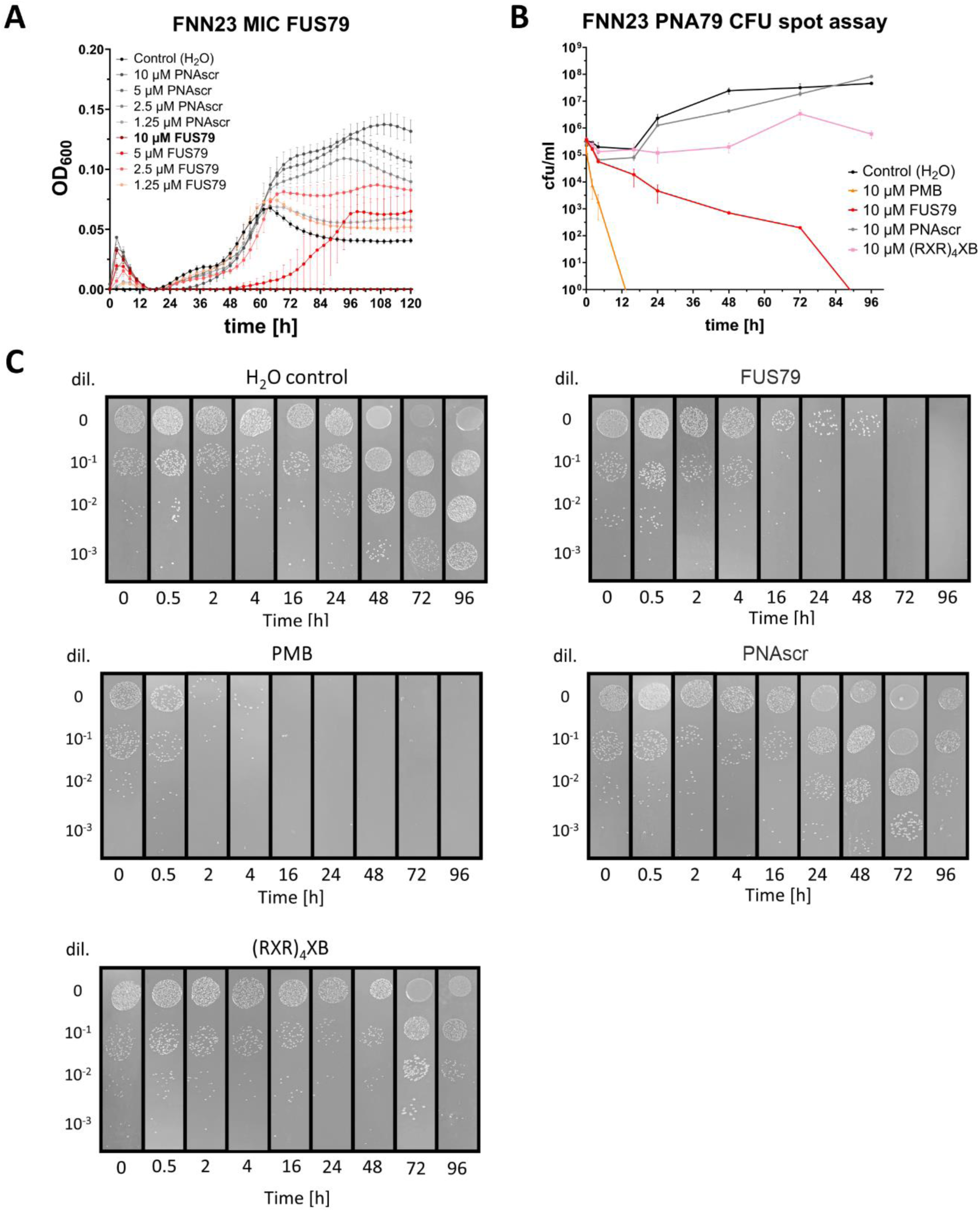
FUS79 is bactericidal against FNN23 at 10 µM. (A) Growth kinetics of FNN23 incubated with FUS79 in MHB at 1 x 10^5^ CFU per ml input. The growth curves are depicted as OD_600nm_ over time for three replicates from different o.n. cultures, error bars indicate standard deviation of three experiments. (B) Determination of bactericidal kinetics for H_2_O as negative control, polymyxin B (PMB) as positive control, 10 µM unconjugated (RXR)_4_XB, the corresponding scrambled control (PNAscr) or FUS79. At selected time points post treatment, 100 µl of each sample were collected and diluted 1:10 in 1x PBS to enumerate the number of viable cells (CFU/ml) via spotting on BHI agar plates. After three days CFUs were counted for three technical replicates and quantified. Error bars represent standard deviation of three experiments. (C) Exemplary images out of three replicates showing CFUs on BHI plates after three days incubation from H_2_O control, PMB, (RXR)_4_XB, PNAscr or FUS79.

To explore if the observed growth inhibition was due to bacteriostatic or bactericidal effects, we incubated FNN23 with 10 µM of FUS79 and performed CFU spot assays to quantify bacterial viability. Incubation with FUS79 reduced the CFU count from 24 hours onwards, with no visible CFUs detected on the agar plate after 96 hours (Fig. 3B and C). Treatment of FNN23 with PNAscr or unconjugated (RXR)_4_XB had no bactericidal effect. Polymyxin B (PMB), known to have strong bactericidal activity, fully diminished CFUs after 96 hours (Fig. 3C). Taken together, our results show that FUS79 completely inhibits the growth of FNN23 at 10 µM.

### FUS79 inhibits growth of various fusobacterial strains but not F. nucleatum ssp. vincentii

To better understand the intriguing bactericidal activity of FUS79, we tested whether the observed bactericidal effect was exclusive to FNN23. We performed growth inhibition assays with five other fusobacterial strains associated with human physiology and pathology (52): *F. nucleatum* ssp. *nucleatum* ATCC 25586 (FNN25), *F. nucleatum* ssp. *animalis* 7_1 (FNA), *F. nucleatum* ssp. *vincentii* ATCC 49256 (FNV), *F. nucleatum* ssp. *polymorphum* ATCC 10953 (FNP), and *F. periodonticum* 2_1_31 (FPE) (Fig. 4A). These strains differ in their growth rate. FNN23 and FNA display a long lag phase and show regrowth in MHB after 24-40 hours with a maximum OD_600nm_ value below 0.1. In contrast, FNN25, FNV, FNP and FPE show steady regrowth around 24 hours and reach higher OD_600nm_ values (Fig. 4B).

**Fig. 4:**
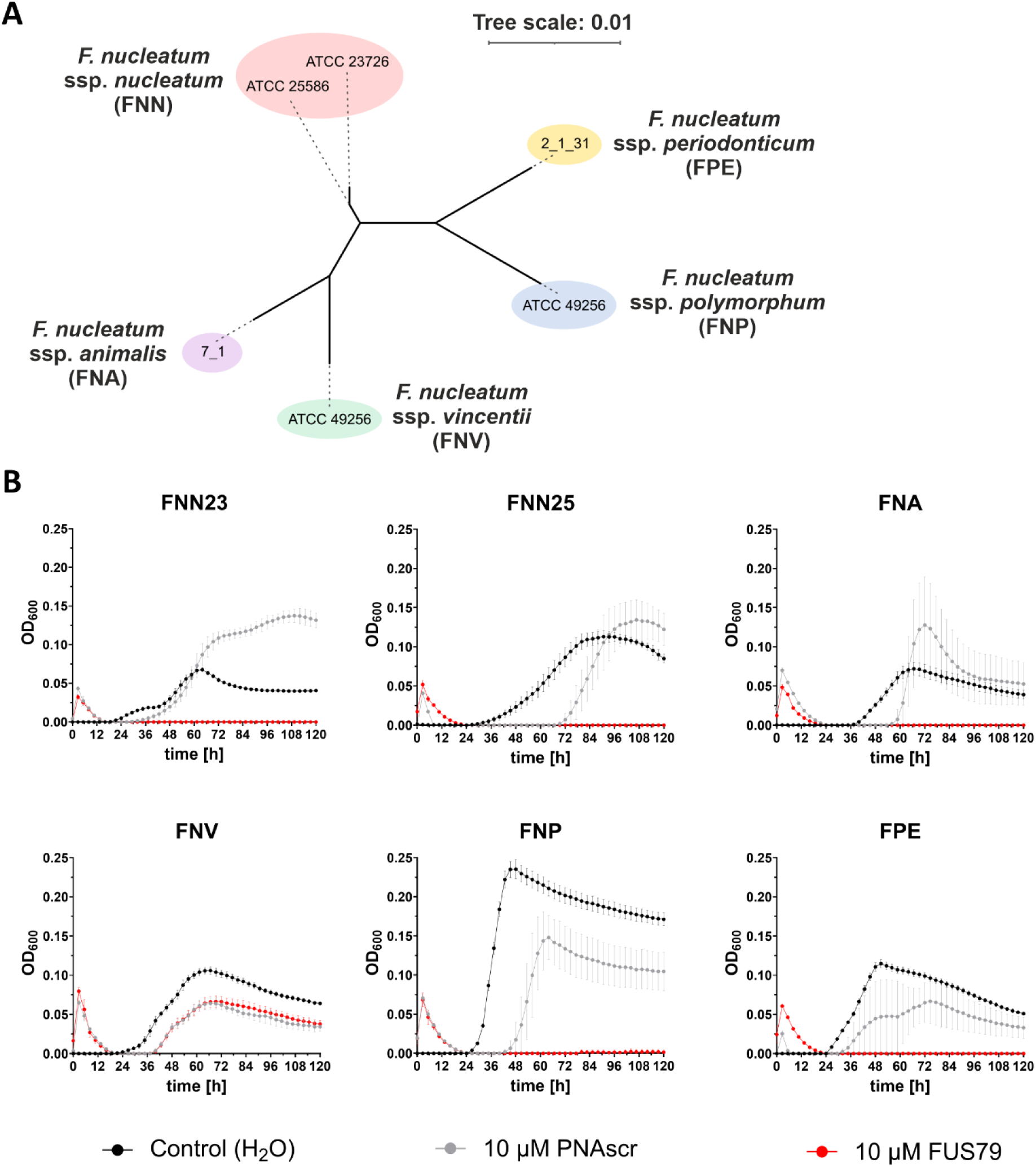
FUS79 inhibits growth of serval fusobacterial species except FNV. (A) Phylogenetic tree of F. *nucleatum* strains and FPE was generated using the respective 16S rRNA sequences. Multiple sequence alignment was performed with MAFFT and a maximum-likelihood phylogenetic tree was then inferred from the aligned 16S rRNA sequences. Branch lengths indicate the expected number of substitutions per site. The scale bar corresponds to 0.01 substitutions per site i.e., one substitution per 100 bases. (B) Growth kinetics of FNN23, FNN25, FNA, FNV, FNP and FPE in MHB at 1 x 10^5^ CFU per ml input incubated with 10 µM FUS79 (red) or the corresponding PNAscr (gray). Growth curves are depicted as OD_600nm_ over time for three replicates from different o.n. cultures, error bars indicate the standard deviation of three experiments.

Remarkably, FUS79 inhibited growth of all strains, except FNV (Fig. 4B). Even though FNV is phylogenetically closer to FNN23 than e.g., FNP or FPE (Fig. 4A), the latter strains are both susceptible to FUS79. When treating the different strains with the PNAscr we observed growth retardation compared to the water control, which might be attributed to the antibacterial activity of (RXR)_4_XB itself. These results indicate that the bactericidal effect of FUS79 is not limited to FNN23, but common among different fusobacterial strains.

### FUS79 does not inhibit growth of several other Gram-negative bacteria

To investigate if FUS79 shows similar bactericidal effects against other bacterial species, we tested its effect on *P. gingivalis*, a Gram-negative anaerobic bacterium that shares the oral niche and co-aggregates with *F. nucleatum* (53). We also included *E. coli* K12 and *E. coli* Nissle 1917 as representative gut commensals (54). Incubation with FUS79 showed no growth inhibition for these species, similar to the PNAscr control (Fig. 5). These data suggest that FUS79 might specifically inhibit certain fusobacterial strains. Furthermore, we tested the effect of FUS79 against *Salmonella enterica* serovar Typhimurium SL1344. We had shown previously that incubation of *S. enterica* with 10 µM (RXR)_4_XB-PNAs causes a growth defect (32), which we confirm here (Fig. 5 bottom right). This suggests that (RXR)_4_XB-PNAs generally impair the growth of *S. enterica* and that FUS79 has no specific bactericidal effect.

**Fig. 5:**
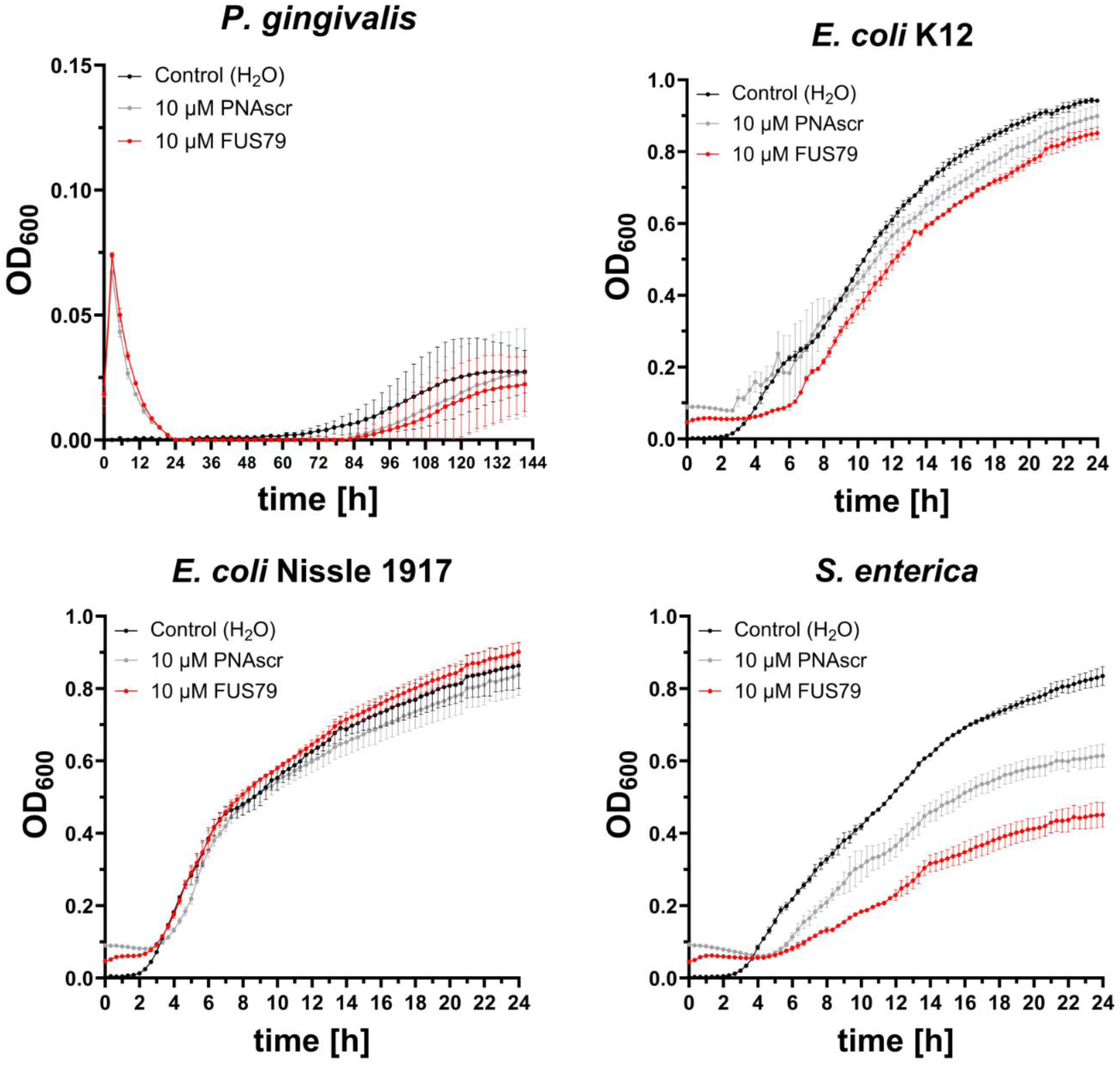
FUS79 does not specifically inhibit growth of other Gram-negative bacteria. Growth kinetics of *P. gingivalis*, *E. coli* K12, *E. coli* Nissle 1917 and *S. enterica* incubated with 10 µM FUS79 or the respective PNAscr control in MHB using 1 x 10^5^ CFU per ml input. For anaerobic growth OD_600nm_ was measured for 144 hours, whereas for aerobic growth data was collected for 24 hours. Growth curves are depicted as OD_600nm_ over time for three replicates from different o.n. cultures, error bars indicate the standard deviation of three experiments.

### The growth inhibitory capacity of FUS79 does not seem to be mediated by off-target regulation

The nucleobase sequence of FUS79 (GACATAATTGT) has no predicted full antisense match within any TIR in FNN23 or FNN25, and no full-length complementary sequences in the genomes of FNA, FNP and FPE. However, we found 19 potential FUS79 TIR off-targets in FNN23 if we allowed one or two mismatches. We reasoned that those potential off-targets are worth investigating since terminal double mismatches in a 10mer PNA do not necessarily abrogate growth-inhibitory activity (45). Since FUS79 showed growth inhibition in five fusobacterial strains, but not in FNV, we searched for mismatch TIR off-targets present in the five vulnerable strains but absent in the resistant FNV. We found one off-target that fits these criteria: FUS79 is complementary to the TIR of *trpB* mRNA (tryptophan synthase subunit beta; C4N14_04955 in FNN23) with two terminal mismatches, i.e., G<u>A</u>CATAATT<u>G</u>T. We tested if a fully complementary PNA sequence targeting *trpB* inhibits the growth of the most vulnerable strain FNN23. Treatment with 10 µM (RXR)_4_XB-*trpB* did not result in growth impairment, arguing against a potential translational inhibition of *trpB* as the mode of action of FUS79 (Fig. S2C).

### Antimicrobial activity of FUS79 is mediated by (RXR)_4_XB and distinct PNA sequence elements

Since FUS79 did not seem to inhibit *F. nucleatum* growth through TIR targeting, we further investigated the potential reasons for the observed toxicity in the five vulnerable strains. To this end, we tested FUS79 without the (RXR)_4_XB module (namely PNA79), with a shortened (RXR)_4_XB module (namely (RXR)_3_XB), coupled to (KFF)_3_K instead of (RXR)_4_XB (Fig. S3A), and two sequence variants of the original FUS79 (Fig. 6A). These PNA sequence variants contain nucleobase substitutions expected to disrupt a predicted 3-bp hairpin structure of FUS79 (Fig. 6A). Even though single-stranded PNAs are presumed to form compact structures due to their flexible backbone and hydrophobicity, they have also been shown to form hairpins under certain conditions (55). After performing growth inhibition assays with FNN23, we observed that only the original FUS79 as well as sequence variant 1 conjugated to full length (RXR)_4_XB inhibited the growth of FNN23 at 10 µM (Fig. 6B). Conjugating PNA79 to (KFF)_3_K did not lead to growth inhibition (Fig. S3A). When testing these compounds in the other fusobacterial strains, only FUS79 and sequence variant 1 were effective in strains susceptible to FUS79 but not in the resistant strain FNV, highlighting a common mode of action between the original FUS79 and variant 1 (Fig. 6C). Therefore, we conclude that intact (RXR)_4_XB together with specific parts of the FUS79 sequence (GACATA<u>W</u>T<u>W</u>GT) appear to be essential for the bactericidal effect.

**Fig. 6:**
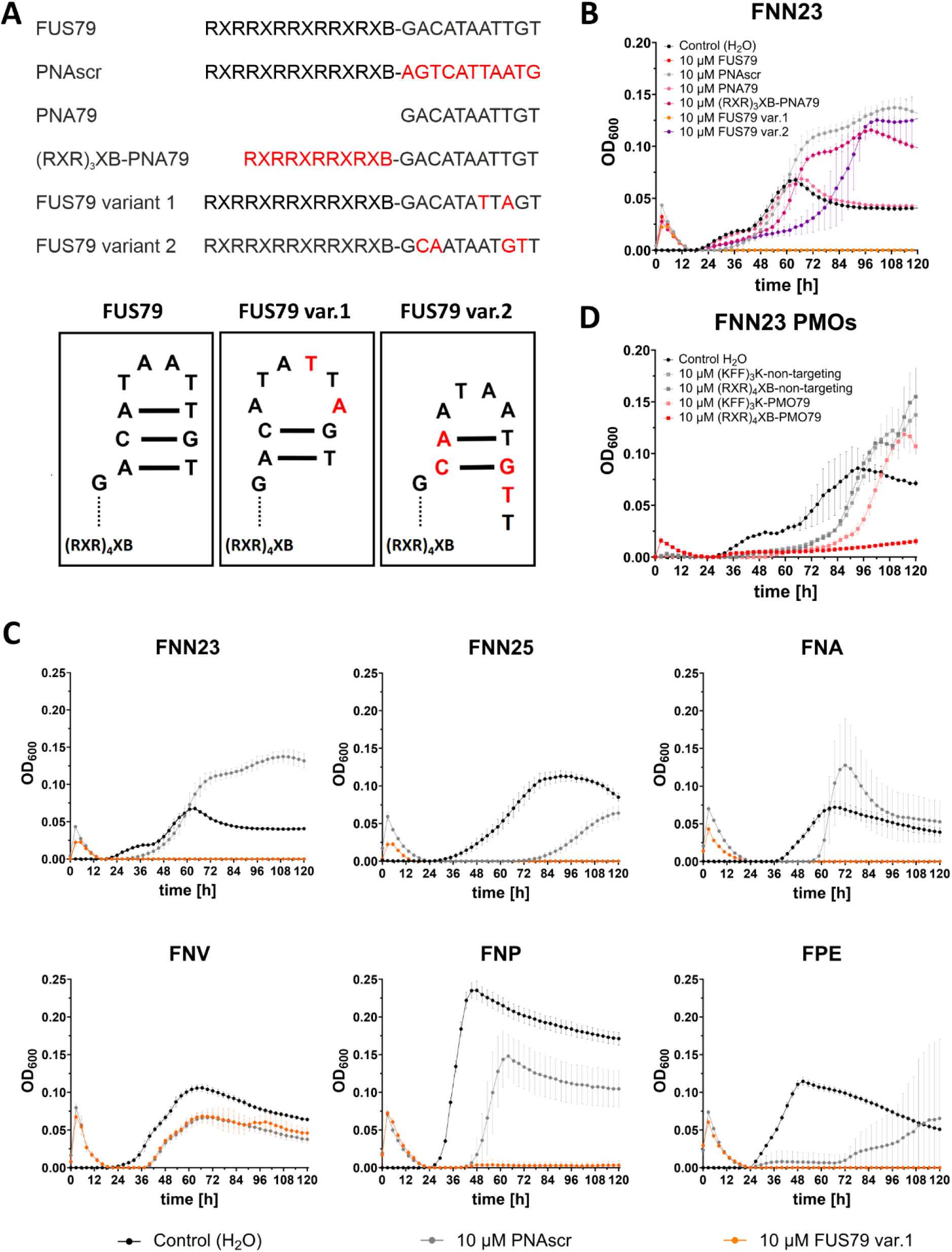
Bactericidal effect of FUS79 is dependent on (RXR)_4_XB and certain sequence elements of PNA79, but independent of ASO modality. (A) (Top) List of investigated FUS79 variants. Changes in respect to the original FUS79 conjugate are depicted in red. (Bottom) Schematic representation of the putative hairpin structure of PNA79. Nucleobase exchanges, shown in red for variant 1 and variant 2, including the presumed changes in structure. (B) Growth kinetics of FNN23 incubated with CPP-PNAs or PNAs at 10 µM concentration. The growth curve of FUS79 (red) is nearly identical to variant 1 (orange) and mostly overlaps with the orange line. (C) Growth kinetics of FNN23, FNN25, FNA, FNV, FNP and FPE incubated with 10 µM FUS79 variant 1 (orange) and corresponding PNAscr (gray). (D) Growth kinetics of FNN23 incubated with 10 µM of (KFF)_3_K- or (RXR)_4_XB-conjugated PMO79 shown in red. Water control is depicted in black, respective non-targeting PMO controls in gray. All growth curves are shown as OD_600nm_ over time for three replicates from different o.n. cultures with 1 x 10^5^ CFU per ml input in MHB, error bars indicate the standard deviation of three experiments.

We also tested different experimental conditions, i.e., higher bacterial inoculum density and alternative bacterial culture media. These experiments showed that at 10 µM, FUS79 is no longer bactericidal at 10^7^ CFU per ml in MHB (Fig. S3B), and has no effect when tested in peptide-rich media such as Brain Heart Infusion (BHI) or Columbia Broth (ColB) (Fig. S3C). This demonstrates the MIC of FUS79 depends on the inoculum size and medium type, which is common in many antibiotics and antimicrobial peptides (56, 57).

To further explore if the antibacterial activity was dependent on the ASO backbone modality, we replaced the PNA moiety with a phosphorodiamidate morpholino oligomer (PMO), while retaining the nucleobase sequence of FUS79. Although antisense-mediated translation inhibition by PMO typically requires a longer sequence compared to PNA (58, 59), we decided to keep the same nucleobase length to directly compare the two conjugates, since the mode of action of FUS79 is unlikely to be mediated by sequence-specific binding. The PMO-ASO was conjugated to either (RXR)_4_XB or (KFF)_3_K as control since (KFF)_3_K-PNA79 had shown no bactericidal effect. At 10 µM concentration, (RXR)_4_XB-PMO79 showed similar growth inhibition as FUS79 even though the backbone of PMO is very different from PNA. In contrast, the (RXR)_4_XB-conjugated non-targeting PMO control as well as (KFF)_3_K-PMO79 did not inhibit growth of FNN23 (Fig. 6D). This result suggests that the bactericidal effect is due to the combination of the ASO nucleobase sequence and (RXR)_4_XB.

### RNA-seq analysis of a sensitive and resistant F. nucleatum strain upon FUS79 treatment

To investigate the mechanism of FUS79, we employed RNA-seq to measure transcriptomic changes associated with treatment in a sensitive fusobacterial strain. We chose FNN23, which was the strain most susceptible to FUS79, as well as FNV, which was resistant. We treated both strains with FUS79, the corresponding PNAscr, or water as control and took RNA-seq data at 30 minutes to monitor early transcriptomic changes prior to the onset of growth inhibition and at 16 hours after the onset of the bactericidal activity (Fig. 7A).

**Fig. 7:**
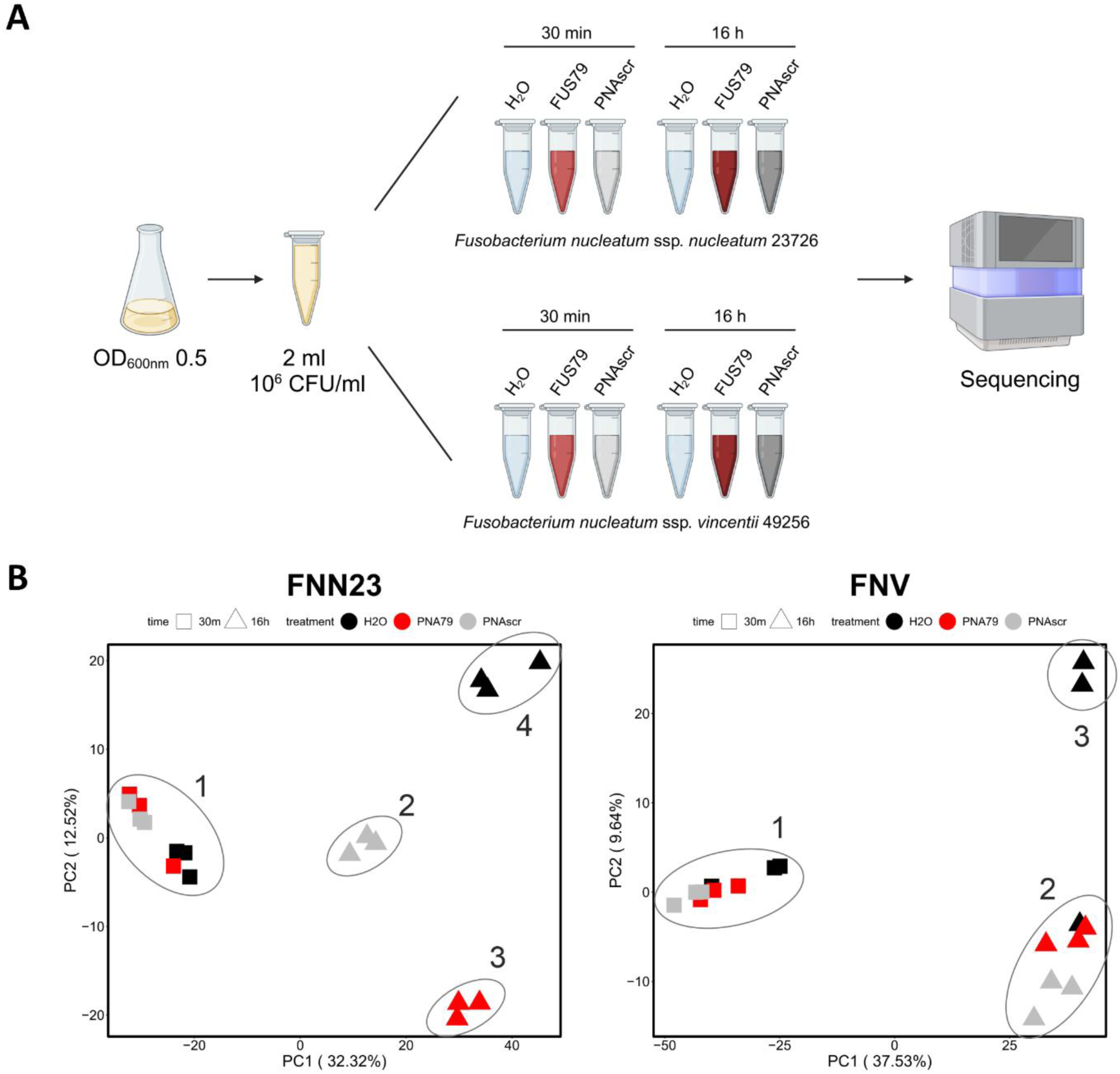
PCA plot reveals distinct clusters for FUS79-treated sensitive FNN23 versus resistant FNV. (A) Experimental workflow showing the different samples subjected to RNA-seq. Image has been created with BioRender.com. (B) Principal component analysis (PCA) of 18 samples for each fusobacterial strain after TMM normalization. Clusters were inserted manually grouping closely positioned samples.

To obtain an overview of the data, we evaluated the results using principal component analysis (PCA; Fig. 7B and Fig. S4). When comparing PC1 vs. PC2 (Fig. 7B) at 30 minutes, there were no large differences between samples for either strain. In contrast, at 16 hours post-treatment, we observed distinct patterns of clustering for FNN23 and FNV, as might be expected given the different sensitivity of these strains to FUS79. In FNN23, the water control, FUS79 and the PNAscr formed three distinct clusters, indicating disparate responses to these three treatment conditions. In contrast, for FNV samples, we observed only two clusters: one containing two of the water-treated control samples, and a second containing both FUS79- and PNAscr-treated samples as well as a single water-treated control, which was removed from further analysis. PC1 vs. PC3 (Fig. S4A) as well as PC2 vs. PC3 (Fig. S4B) showed a similar clustering and did not further help to separate the samples. These results indicate that a 16 hour FUS79 treatment elicits a distinct response from the PNAscr in strain FNN23, which is sensitive to the ASO, but not in the resistant strain FNV.

To investigate general transcriptomic responses of fusobacteria to CPP-PNA treatment, we compared FUS79 as well as PNAscr treated samples to the water control (Supplementary data SD1). We mapped the reads using the *F. nucleatum* ssp. *nucleatum* ATCC 23726 annotation file (NZ_CP028109.1) with custom annotation and the *F. nucleatum* ssp. *vincentii* 3_1_36A2 annotation file (NZ_ CP003700.1). We observed that while FUS79 and PNAscr do not induce significant changes (log_2_ fold change <−1.8 or >1.8 with an adjusted false discovery rate of ≤ 0.01) after 30 minutes in FNN23, both induce downregulation of transcripts involved in purine metabolism after 16 hours (C4N14_07225-C4N14_07265, Fig. S5). Upregulated transcripts differ strongly between the two CPP-PNA treatment conditions (Fig. S5). For FNV, only *hemC*, which is important for heme biosynthesis, is upregulated upon PNAscr treatment after 30 minutes. At 16 hours post treatment, both CPP-PNAs downregulated *pckA* and *pdxS*, involved in gluconeogenesis and cofactor biosynthesis, respectively. Furthermore, mRNAs HMPREF0946_RS04730, HMPREF0946_RS03355 and HMPREF0946_RS08775 as well as sRNA FunR47 are downregulated, but there are no commonly upregulated transcripts (Fig. S5). In conclusion, there does not appear to be a common CPP-PNA-induced transcriptomic response in fusobacteria.

### FUS79 induces the σE membrane stress response in the sensitive F. nucleatum strain

To further characterize the distinct transcriptional response to FUS79, we focused on the differences between FUS79 and PNAscr (Fig. 8), though we have made results for all relevant comparisons available (Fig. S5). In keeping with our PCA analysis, we see minimal differences in RNA levels in both FNN23 and FNV after 30 minute treatment with either of the two CPP-PNAs (Fig. 8A and 8B). However, at 16 hours following CPP-PNA treatment we observe pronounced differences between FUS79 and PNAscr in FNN23 samples. Specifically, we found a total of 82 differentially expressed transcripts between FUS79 and PNAscr samples (Fig. 8A). Interestingly, six of the top ten upregulated transcripts upon FUS79 treatment compared to PNAscr belong to the σE regulon of FNN23 (60) (marked with red boxes in Fig. 8A). The σE regulon plays an important role in the global stress response in Gram-negative bacteria and is composed of multiple proteins and small RNAs (sRNAs) involved in the maintenance of envelope integrity (61, 62). Recently, the σE regulon has been studied in *F. nucleatum* showing an oxygen induced stress response reminiscent of the activated σE response in Proteobacteria (60). Indeed, closer inspection of the transcriptome after 30 minutes treatment shows that the most highly upregulated gene is σE itself (*rpoE*, C4N14_03400; indicated in Fig. 8A left panel).

**Fig. 8:**
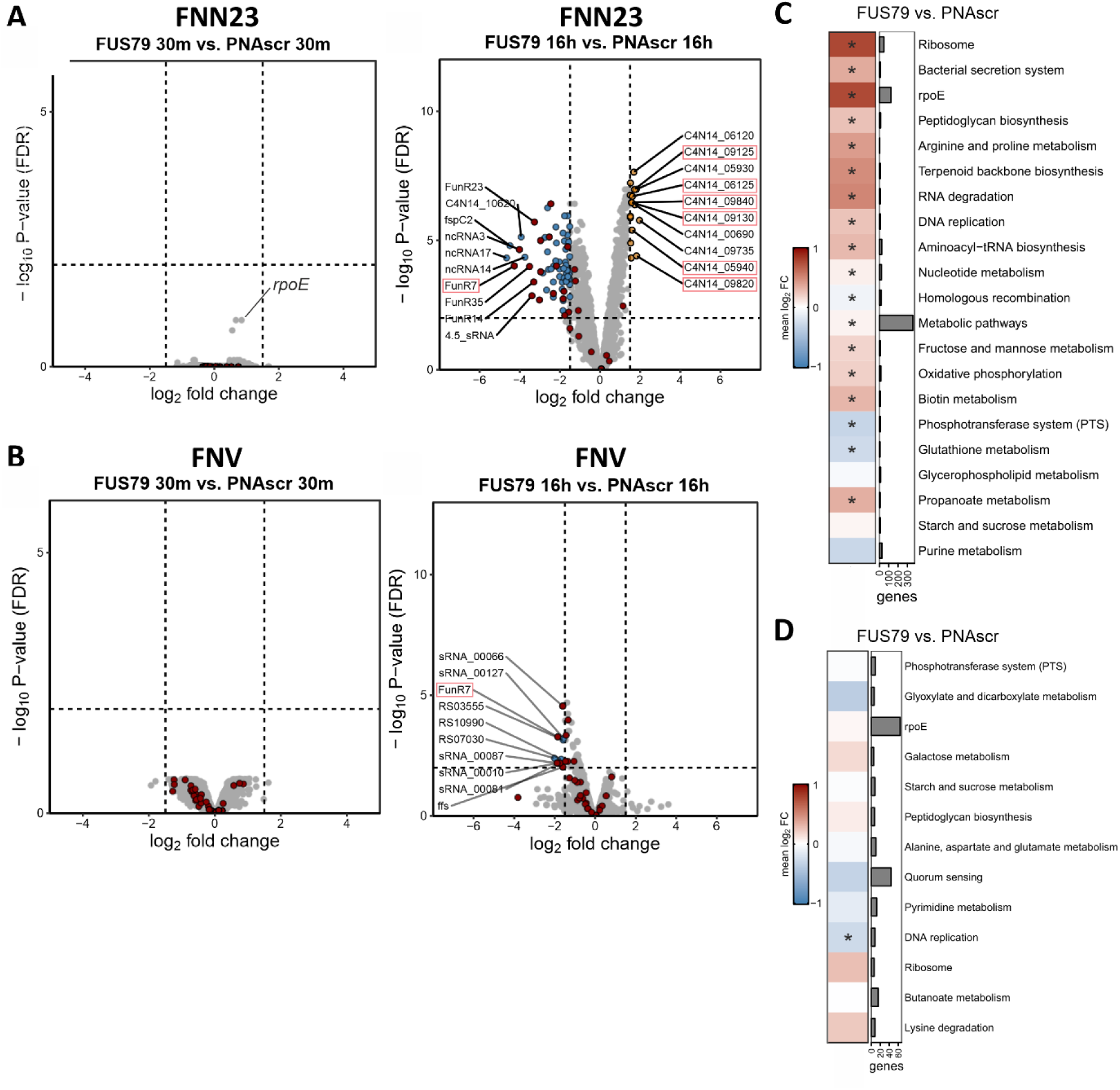
FUS79 treatment induces σE-dependent membrane stress response in sensitive FNN23. (A, B) Transcriptomic response of FNN23 (A) and FNV (B) upon FUS79 or PNAscr treatment. Volcano plots show differential gene expression as –log_10_ false discovery rate (FDR)-adjusted P-values on y-axis and log_2_ fold change on x-axis. (Left) 30 minutes 10 µM FUS79 vs. 10 µM PNAscr. (Right) 16 hours 10 µM FUS79 vs. 10 µM PNAscr. Significantly differentially expressed transcripts are defined by a log_2_ fold changes <-1.8 or >1.8 and an FDR adjusted P-value <0.01, depicted as dashed lines in the plot. Significantly upregulated transcripts are depicted as orange dots, significantly downregulated transcripts as blue dots. All sRNAs are colored as red dots. The top ten differentially expressed transcripts are specified by the indicated locus tag. Gene locus tags that are part of the σE (*rpoE*) regulon are red-boxed. (C, D) KEGG pathway enrichment analysis of annotated gene sets and the manually added σE regulon for FNN23 (C) or FNV (D) after 16 hours treatment. RNA-seq data of FNV and FNN23 was analyzed using annotated KEGG pathways. (*) FDR-adjusted P-value adjusted <0.001. Gene number per pathway is indicated on the right and all pathways are shown as median log_2_ fold change with the corresponding color shade as indicated in the legend on the left.

The function of σE was shown to be supported by two sRNAs, namely FoxI and FoxJ, which regulate mRNAs of envelope proteins (44). However, neither FoxI nor FoxJ were differentially expressed in FNN23. Of all the differentially expressed sRNAs, many were downregulated upon FUS79 treatment, but none upregulated (red dots, Fig. 8A). Out of the sRNAs that are part of the σE regulon, only FunR7 was significantly downregulated. Further, the 4.5S signal recognition particle RNA (4.5S RNA), which is generally involved in bacterial protein synthesis as well as in directing the translation of membrane and secretory proteins to the inner membrane (63, 64), is among the top ten downregulated transcripts. The two strongest downregulated mRNAs are C4N14_10620, a hypothetical protein and *fspC2*, a small ORF uncharacterized in fusobacteria. Collectively, our RNA-seq analysis shows upregulation of transcripts involved in the membrane stress response of FNN23 upon FUS79 treatment, with particular upregulation of transcripts in the σE regulon.

The resistant strain FNV displays no significantly upregulated transcripts upon FUS79 treatment compared to the PNAscr after 16 hours (Fig. 8B). Six of the ten most downregulated transcripts are sRNAs including FunR7 and the 4.5S RNA (ffs), possibly indicating a partly common response to FUS79 shared between both strains. Apart from the shared downregulation of FunR7 and 4.5S RNA, the transcriptomic response to FUS79 versus PNAscr has no other common aspects between the vulnerable FNN23 and the insensitive FNV strain. In conclusion, we observed that the transcriptomic response between FNN23 and FNV are distinct and only the susceptible strain shows upregulation of the σE regulon.

### FUS79 induces general stress responses in FNN23 but not in FNV

To put our differential expression analysis in the context of functional pathways, we performed a gene set enrichment analysis using the KEGG database supplemented with the σE regulon of *F. nucleatum*. We found that upon FUS79 treatment there was a strong induction of KEGG pathways associated with ribosomes, arginine and proline metabolism, bacterial secretion systems, peptidoglycan synthesis, RNA degradation and terpenoid backbone synthesis in the vulnerable strain FNN23 (Fig. 8C). Many of these pathways are linked to a general stress response, response to antibiotic treatment or membrane homeostasis (65–67). As seen above, the σE regulon was strongly upregulated as well. Two pathways associated with the phosphotransferase system and glutathione metabolism were downregulated. These pathways are mainly involved in carbohydrate transport, transcription regulation, stress response and the regulation of cell division (68–70). In contrast, analysis of the resistant FNV strain revealed no pathways that were significantly upregulated and only the DNA replication pathway to be significantly downregulated (Fig. 8D).

Taken together, our RNA-seq analysis revealed a strong transcriptomic response characterized by the upregulation of the σE regulon, antibiotic stress responses and membrane homeostasis after 16 hours for FUS79 compared to PNAscr in FNN23 but not in the resistant strain FNV.

## DISCUSSION

In our attempt to design species-specific asobiotics targeting *F. nucleatum*, we found that although (RXR)_4_XB was able to efficiently penetrate FNN23, none of the CPP-ASOs that we designed to target putative essential mRNAs inhibited bacterial growth. Instead, we observed an intriguing bactericidal effect of a PNA designed as non-targeting control against five fusobacterial strains. Below we propose potential reasons for our inability to establish target-specific antibacterial ASOs in fusobacteria, such as poor cytosolic delivery of CPP-PNA conjugates, limited knowledge of gene essentiality as well as PNA design. Further, we discuss potential mechanisms of FUS79 and why FNV might be resistant. Overall, our observations highlight the importance of considering and reporting unanticipated antibacterial activities, which is essential when evaluating CPP-ASOs as tools for precision microbiome editing (20).

### Exogenously delivered ASOs show no sequence-dependent antisense activity in F. nucleatum

Serval arguments support the idea that PNAs might function effectively in fusobacteria. First, *F. nucleatum* possesses of a large number of regulatory sRNAs (60, 71, 44). Inferring from Gram-negative model species of bacterial RNA biology (72, 73), the majority of these fusobacterial sRNAs can be assumed to repress or activate targets mRNAs by targeting their 5’ end. Indeed, we have already shown that the *F. nucleatum* sRNA FoxI represses *fomA* mRNA by base pairing around the start codon (60). Thus, fusobacterial mRNAs are permissive to antisense modulation. Second, in line with previous reports of arginine-rich peptides as efficient ASO delivery agents in serval Gram-negative species (74), we demonstrated that fluorophore-labeled (RXR)_4_XB efficiently penetrates the membrane of FNN23. Third, we screened many genes, which are essential and shown to be amenable to PNA-mediated inhibition in bacteria such as *E. coli* (25, 75), *S. enterica* (76), *Staphylococcus aureus* (77), *K. pneumonia* (74), and *Buchnera aphidicola* (78).

Even though out of the ASO targets we tested, only *ftsZ* is experimentally confirmed to be essential in FNN23 (46), we expect the other targets to encode essential gene products as well based on observations from other bacteria (79, 80). That said, despite initial transposon screens and the recent establishment of CRISPRi for FNN23 and FPE (46, 79, 81, 82), we lack a complete picture of gene essentiality in fusobacteria. It is also possible that the designed PNA sequences are not efficient enough to induce growth inhibition. The AT-rich genome of fusobacteria makes it difficult to design PNAs with a high melting temperature and low self-complementarity (83, 84), both important factors for ASO efficiency (47, 85). Moreover, the exact binding site on the mRNA target also greatly influences the translational inhibitory capacity of an ASO (86). It might be that we did not yet identify the best target region to inhibit mRNA translation with ASOs in fusobacteria. A sequence tiling screen within TIRs might help to identify the most vulnerable binding window to increase ASO efficiency. However, slow and fastidious growth of fusobacteria complicates the establishment of robust growth assays with a clear separation of ASO-triggered growth inhibition versus normal growth fluctuations.

Although (RXR)_4_XB was able to deliver TAMRA into fusobacteria, it might not effectively deliver the much larger PNAs to the bacterial cytoplasm. Indeed, the delivery efficiency of CPPs was shown to depend greatly on the chemical properties of the cargo (24, 87). It is therefore worth considering testing alternative ASO delivery vehicles for *F. nucleatum*. Gold nanoparticles, vitamin B12 and DNA-tetrahedrons have been reported to facilitate ASO uptake by Gram-negative and Gram-positive bacteria (84, 88–90). Another promising delivery approach are siderophores, because these small Fe(III) chelating molecules produced by bacteria (91) are not cytotoxic. Indeed, siderophores have been employed in a “Trojan horse” strategy to delivery antibiotics into bacteria (92). They have also demonstrated good delivery efficacy of PNAs and PMOs into *E. coli* (93, 94). Although fusobacteria lack confirmed siderophore biosynthesis genes, they might be able to utilize xenosiderophores, i.e., iron-chelating compounds produced by other members of the microbiota as shown for *Clostridioides difficile* (95). It remains to be seen which other delivery mechanisms are feasible for ASO delivery into *F. nucleatum*.

Overall, our results highlight that although asobiotics represent a promising antibacterial strategy for multiple Gram-negative species, application to non-model bacteria faces challenges highlighting the need for further studies to systematically investigate ASO delivery and application in non-model bacteria.

### Bactericidal effect of FUS79

We have identified a non-targeting (RXR)_4_XB-PNA (FUS79), which is bactericidal against serval fusobacterial strains. It seems unlikely that FUS79 inhibits growth of sensitive strains via an antisense off-target mechanism. First, FUS79 has no fully complementary sequence within any TIR in FNN23. Second, the 19 potential FNN23 TIR off-targets with one or two terminal mismatches as well as transcriptome-wide off-targets were not significantly deregulated after FUS79 treatment in our RNA-seq experiments. As some bacterial sRNAs were shown to prefer imperfect matches (96), we investigated the transcript levels of the mismatched TIR off-target *trpB* (C4N14_04955). However, also *trpB* did not show significant mRNA downregulation upon FUS79 treatment compared to the PNAscr control (Supplementary data SD1). Third, FUS79 has no fully complementary sequence in any genomic region of FNA, FNP or FPE. Fourth, the FUS79 sequence variant 1, which has two nucleobase substitutions at non-terminal positions showed the same antibacterial effect as FUS79, although the resulting mismatches should hamper binding to a FUS79 complementary sequence. Nevertheless, we cannot exclude that mismatched off-targets might lead to translational inhibition without affecting mRNA abundance. In that regard it was shown that target mRNA depletion is not a universal trait of PNA-mediated translation inhibition (25).

Our RNA-seq profiles of FNN23 bacteria treated with FUS79 show upregulation in transcripts involved in membrane homeostasis. The bactericidal effect of FUS79 might therefore be mediated by membrane damage triggered by the specific CPP and ASO sequence combination. The resistance of FNV could be mediated by lipopolysaccharide (LPS) modifications affecting CPP entry. Based on genomic analysis, the LPS O-antigen of FNV is different compared to the sensitive FNN25. FNV possesses genes necessary for the incorporation of sialic acid, galactopyranose, galacturonate and colitose into LPS, whereas other *F. nucleatum* ssp. *nucleatum* strains do not (97). Indeed, changes in the LPS composition of *E. coli* were shown to affect the uptake of CPP-PNA conjugates (98). That said, the sensitive FNP also incorporates sialic acid into its O-antigen (99) and FNV (RXR)_4_XB-TAMRA uptake rates are similar to the sensitive FNN23 (Fig. S6).

### FUS79 is a strong inducer of the fusobacterial σE response

We found that FUS79 induces many transcripts involved in membrane biosynthesis and homeostasis in the vulnerable FNN23 strain. KEGG pathway analysis further confirmed that the σE regulon was strongly induced. It is worth noting that we added this gene group manually because recent studies showed that more genes are part of the σE regulon than computationally predicted in the KEGG database (60, 44). An enrichment in this regulon could therefore also result from the high number of individual upregulated transcripts present in multiple pathways and not be strictly σE-specific. Nevertheless, the upregulation of *rpoE* in response to FUS79 after 30 minutes indicates an induction of the σE stress response. The σE regulon is activated via numerous stressors such as heat shock, osmotic stress, pH stress, oxidative stress or antibacterial agents (100, 101, 61, 102). We cannot link the activation of σE to membrane stress specifically as the KEGG analysis demonstrated the activation of diverse stress responses. In FNN23, activation was recently shown to be induced upon oxygen exposure rather than membrane stressors such as polymyxin B or lysozyme (60). However, these experiments were conducted in cation- and peptide-rich ColB, where bacteria are generally less sensitive to antimicrobial compounds like CPPs (103, 104) and FUS79 did not retain its bactericidal effect. In contrast, in this study we used MHB, which might lead to an overall higher sensitivity to membrane targeting antimicrobials.

We propose that the unique combination of (RXR)_4_XB and the PNA79 sequence might interact with the bacterial envelope and lead to σE regulon activation as well as a membrane stress response with subsequent growth inhibition upon membrane disruption. Further mechanistic studies are needed to determine the precise mode of action of FUS79. The MIC of FUS79 depended on the medium type and inoculum size, as is often described for membrane-acting antimicrobial agents (56, 105, 106). Moreover, a (KFF)_3_K-conjugated PNA79/PMO79 did not impair bacterial growth (Fig. S3A and Fig. 6D), supporting the hypothesis that the cytotoxicity is dependent on (RXR)_4_XB and certain sequence elements. It would be interesting to test if a CPP with a similar uptake efficiency and entry mode as (RXR)_4_XB would display bactericidal activity when coupled to PNA79. This would test whether (RXR)_4_XB is specifically needed for the toxicity of FUS79 or whether it depends on the cellular delivery of the ASO component. In general, the conjugation of a CPP to an ASO changes the overall chemical properties as well as the secondary structure of both components, which might influence the cell-penetrating and target binding efficiency of the conjugate (105, 107, 108). While the net positive charge of (RXR)_4_XB should be unaffected by conjugation to the charge neutral PNA or PMO modalities (41, 109), changes in the CPP conformation are more likely. These changes might lead to increased penetration as membrane disruption was shown to be dependent of the CPP structure in Gram-negative bacteria (110).

### Outlook

Broad-spectrum antibiotics can deplete *F. nucleatum* from the tumor site leading to therapeutic benefit, but such treatment would disrupt the patient’s protective microbiota. Species-specific ASOs could offer a solution to this problem. We tried to establish asobiotics against *F. nucleatum*, but were not able to achieve antisense-mediated growth inhibition. However, it remains unclear whether the CPP-PNAs effectively reach the fusobacterial cytoplasm and are able to bind their mRNA target. Mass spectrometry of cellular fractions could be a feasible approach to test CPP-PNA localization, but our preliminary experiments were unsuccessful. As discussed above, further systematic analyses are required to explore asobiotics as antimicrobial agents against *F. nucleatum*.

Additional therapeutic alternatives to deplete *F. nucleatum* could be species-specific lytic phages, such as FNU1 (111), narrow spectrum antibiotics (112), targeting the bacterium through chemically modified transfer RNA fragments (113), tailored CRISPR antimicrobials (114) or the administration of monoclonal antibodies targeting fusobacterial proteins (115, 116). FUS79 might also be considered a lead for the development of a selective inhibitor while sparing resident microbiota. Interestingly, FUS79 showed bactericidal activity against several fusobacterial strains, with no effect on *P. gingivalis* or *E. coli* strains. However, FUS79 exhibits an unidentified bactericidal mechanism that seemingly operates independently of target inhibition. We have tried coupling FUS79 to fluorophores or biotin, but the conjugations abolished the bactericidal effect, which limits the interpretation of localization with these modified conjugates.

Identifying the molecular mechanism of FUS79 will be essential for its clinical application. The membrane interaction of FUS79 with *F. nucleatum* could be mechanistically investigated using different approaches such as membrane model systems, membrane dyes or molecular dynamics simulations. Such assays bear challenges, since the envelope structure of *F. nucleatum* is largely unknown and the exposition to atmospheric oxygen triggers membrane stress response itself (60). However, by leveraging the power of medicinal chemistry with respect to chemical modalities and modifications, it should be possible to further enhance the potency of FUS79 to fully harness its potential as a specific antibacterial agent.

## MATERIAL AND METHODS

### Strains and growth conditions

The following fusobacterial strains were used in this study: *Fusobacterium nucleatum* ssp. *nucleatum* ATCC 23726 (FNN23) acquired from the American Type Culture Collection (ATCC), *F. nucleatum* ssp. *nucleatum* ATCC 25586 (FNN25), *F. nucleatum* ssp. *vincentii* ATCC 49256 (FNV) and *F. nucleatum* ssp. *polymorphum* ATCC 10953 (FNP) acquired from the German Collection of Microorganisms and Cell Culture (DSMZ), *F. nucleatum* ssp. *animalis* 7_1 (FNA) and *F. periodonticum* 2_1_31 (FPE) both received as a kind gift from E. Allen-Vercoe (University of Guelph, Canada). All strains were routinely grown at 37°C in 80:10:10 (N_2_:H_2_:CO_2_) anaerobic conditions on 2% agar supplemented BHI plates (brain–heart infusion (BHI), 1% (w:v) dried yeast extract, 1% (v:v) 50% sterile filtered glucose solution, 5 µg per ml of hemin; 1% (v:v) fetal bovine serum) from frozen 20% glycerol stocks kept at -80°C. For liquid growth the strains were cultured in Columbia broth (ColB, BD Difco™). Precultures were prepared 24 h before inoculating the working culture at a 1:50 dilution in fresh ColB until OD_600nm_ 0.5 was reached. Cultures were then diluted in non-cation adjusted Mueller-Hinton Broth (MHB, BD Difco™) to the desired CFU per ml number. *Porphyromonas gingivalis* (DSM 20709) was purchased from DSMZ and grown at 37°C in anaerobic conditions as mentioned above on 2% agar supplemented BHI+ plates (BHI, 1% (w:v) dried yeast extract, 1% (v:v) 50% sterile filtered glucose solution, 5 µg per ml of hemin; 10% (v:v) fetal bovine serum and 1 µg per ml vitamin K3) for four days. For liquid growth *P. gingivalis* was cultured in BHI+ medium (BHI, BD Difco™ supplemented with 1% (v:v) 50% sterile filtered glucose solution, 5 µg per ml of hemin; 1% (v:v) fetal bovine serum and 1 µg per ml vitamin K3). Precultures were prepared 24 h before inoculating the working culture at a 1:20 dilution in BHI+ until OD_600nm_ of 0.5 was reached. All plates, media, buffers and reagents were brought into the anaerobic chamber the day before usage to ensure full oxygen depletion.

Aerobic bacterial culture was conducted with *Escherichia coli* K12 MG1655 (provided by D. Lee), *E. coli* Nissle 1917 and *Salmonella enterica* serovar Typhimurium SL1344 (provided by D. Bumann, Biocenter Basel, Switzerland;). The strains were streaked on lysogeny broth plates, incubated o.n. at 37°C and cultured in non-cation adjusted MHB with aeration at 37°C and 220 rpm constant shaking.

### Cell-penetrating peptides (CPPs) and peptide nucleic acids (PNAs)

CPPs, PNAs and CPP-PNA constructs were obtained from Peps4LS GmbH, where all compounds were tested with mass spectrometry and HPLC to assess their quality and quantity. PMOs were purchased from GeneTools, LLC. In house CPP-PNA synthesis was performed by W. Tegge and B. Kornak at the Helmholtz Centre for Infection Research (HZI, Braunschweig). PNA sequences were designed with the help of MASON (45) and PNA scrambled sequences were verified to have no off-targets in translation initiation regions by manual searches in the whole genome sequence of FNN23 (NZ_CP028109). Comparisons with the other fusobacterial strains were also conducted manually using the genome files NZ_AE009951.2 for FNN25, NZ_CP007062.1 for FNA, NZ_CP003700.1 for FNV, NZ_CM000440.1 for FNP, and NZ_CP028108.1 for FPE. To ensure solubility and correct stock preparation, all compounds were briefly vortexed for three seconds, spun down, heated at 55°C for five minutes and then again vortexed and centrifuged. 200 µM stocks were prepared in water, concentration was determined via Nano-Drop spectrophotometer measurements at A_205nm_ for CPPs or A_260nm_ for PNAs and adjusted if necessary. CPP and PNA stocks were stored at -20°C, low binding tips as well as low binding tubes (Sarstedt) were used throughout for handling.

### Confocal Laser Scanning Microscopy (CLSM) for investigation of CPP uptake

To investigate the efficiency of different cell-penetrating peptides, we excluded a flow cytometry approach to quantify penetration efficiency, because upon cell fixation, CPPs were reported to bind to cell membranes and remain attached even after multiple washes, which would result in a false positive penetration signal in flow cytometry (117). Thus, we opted for a microscopy based investigation with lower throughput but discrimination potential between membrane associated and cytosolic signals. CPPs were coupled to the fluorophore 5(6)-carboxytetramethylrhodamine (TAMRA) by Peps4LS GmbH and 200 µM stocks were prepared in house. Briefly a bacterial overnight culture was diluted to 10^8^ CFU per ml in fresh non-cation adjusted MHB (BD Difco™, Thermo Fisher Scientific), incubated with 5 µM CPP-TAMRA at 37°C with 230 rpm in the anaerobic chamber, taken out of the chamber, centrifuged at 4°C for ten minutes with 13,000 g to collect the pellet, fixated with 4% (w:v) PFA at 4°C for ten minutes, stained with the membrane dye N-(3-Triethylammoniopropyl)-4-(6-(4-(diethylamino)phenyl) hexatrienyl)pyridinium dibromide (FM 4-64 Biomol, 1:1000 diluted in water) at RT for 15 minutes, washed once with 1x PBS and 1-2 μl of cell suspension spotted on an 1.5% agar pad. Samples were imaged using ibdi chambers on a Leica SP5 laser scanning confocal microscope (Leica Microsystems) at the corresponding wave lengths. The emission was detected at 685-795 nm for FM 4-64 and 570-620 nm for TAMRA. CLSM images were analyzed using ImageJ. To quantify CPP uptake 100 cells were chosen out of four images per condition using the bright field image and analyzed for cytosolic TAMRA signal by excluding TAMRA negative cells and TAMRA signal co-localized with the FM 4-64 signal.

### Minimum inhibitory concentration (MIC) determination and growth inhibition assays

To determine the MIC value, broth microdilutions were performed according to the standard protocol with some modifications (118). An overnight bacterial culture was diluted in fresh ColB medium and grown to OD_600nm_ 0.5 (mid-exponential phase). Fusobacteria were then diluted 1:2500 or 1:250 in fresh MHB to obtain 1 x 10^5^ CFU per ml or 1 x 10^6^ CFU per ml as indicated. To determine growth inhibition of *P. gingivalis,* an o.n. culture grown in BHI+ was diluted 1:1500 in MHB to obtain 1 x 10^5^ CFU per ml. *E. coli* K12, *E. coli* Nissle 1917 and *S. enterica* o.n. cultures grown in MHB were diluted 1:1800 in fresh MHB to obtain 1 x 10^5^ CFU per ml. 190 μl of diluted bacterial cultures were pipetted into a transparent 96-well plate (Nunc™, Thermo Fischer Scientific) together with 10 μl of respective 20x CPP or CPP-PNA stock or water as control. Bacterial growth was monitored via measuring OD_600nm_ every 20 minutes with constant shaking in a plate reader (Biotek) positioned in the anaerobic chamber or under normal atmospheric environment with double orbital shaking every 20 minutes prior to each measurment with 237 cpm at 37°C for 120 h. The MIC was defined at the lowest concentration in which growth was visibly inhibited (OD_600nm_ <0.01 for anaerobic cultures and <0.1 for aerobic cultures).

### Determination of bactericidal effect using spotting

To investigate bactericidal effects of PNAs an o.n. culture of FNN23 grown in ColB was diluted 1:50 in fresh ColB and grown to OD_600nm_ 0.5. This culture was then diluted 1:2500 to 1 x 10^5^ CFU per ml in MHB. 190 μl of the culture were dispensed into a transparent 96-well plate (Nunc™, Thermo Fischer Scientific) together with 10 μl of respective 20x PNA stock or water and incubated at 37°C with 237 cpm shaking every 20 minutes. At the respective time points 100 μl were taken out of the well, a 1:10 serial dilution series was prepared with 900 μl anoxic 1x PBS and 5 μl of each dilution were spotted on BHI plates for CFU determination. Plates were incubated for three days at 37°C before taking out the plates of the anaerobic chamber for CFU counting.

### Phylogenetic tree construction

The phylogenetic tree in Figure 4A was built based on 16S rRNA sequences of the six Fusobacterium strains. First, the genomes were screened for 16S rRNA genes using barrnap (v0.9), and the resulting coordinates were used to extract the corresponding sequences with bedtools (v2.31.1). Multiple sequence alignment was performed with MAFFT (v7.490) using default parameters. A maximum-likelihood phylogenetic tree was then inferred from the aligned 16S rRNA sequences using FastTree (v2.1), applying its nucleotide mode and default evolutionary model. The resulting tree was then visualized with iTOL (v7.0).

### PNA treatment for RNA-seq analysis

Three biological replicates for each sample were grown overnight in ColB, diluted the following morning 1:50 in fresh ColB, and after the culture reached OD_600nm_ 0.5, it was diluted 1:250 to 1x10^6^ CFU per ml in MHB. 5 ml of the solution were transferred into 5 ml low-binding tubes (LABsolute) and incubated with 10 μM PNA or the respective volume of water for 30 minutes and 16 hours at 37°C in the anaerobic chamber. The reaction was stopped at the mentioned time points by adding RNAprotect Bacteria (Qiagen).

### RNA isolation

To isolate fusobacterial RNA, samples were incubated 1:1 (v:v) with RNAprotect Bacteria (Qiagen) for five minutes at RT, snap frozen in liquid nitrogen and stored at - 80°C until further processing. To purify RNA the miRNeasy Micro kit (Qiagen) was used. Briefly, cell dilutions were thawed on ice, centrifuged at 4°C with 13,000 g for 20 minutes, the pellets resuspended in 100 μl of 0.5 mg per ml lysozyme (Roth) in TE buffer pH 8.0 and incubated for two minutes at RT. The samples were then incubated with 700 μl Qiazol (Qiagen) for five minutes at RT and after addition of 160 μl chloroform shaken vigorously for 15 seconds to obtain phase separation. The upper aqueous phase was mixed 1:1.5 with 100% ethanol and loaded onto the Qiagen columns. Columns washes, DNase I digest, and elution were performed according to the manual instructions. RNA concentrations were determined via NanoDrop measurements and samples were stored at -80°C until sequencing.

### RNA-sequencing, quantification and differential expression

RNA samples were processed and sequenced at the Core Unit SysMed (University of Würzburg, Germany). RNA quality was investigated using the 2100 Bioanalyzer together with the RNA 6000 Pico kit (Agilent Technologies). Risbosomal RNA (rRNA) depletion was performing using the RiboCop META rRNA depletion kit (Lexogen) according to the manufacturer’s instructions. Depleted RNA was further fragmented via ultrasound for 30 seconds at 4°C. After adapter ligation to the 3’ end, the first-strand cDNA was synthetized using the M-MLV reverse transcriptase. Following purification, 5’ Illumina TruSeq adapters were ligated to the cDNA, the cDNA amplified by PCR and the resulting product purified with the Agencourt AMPure XP kit (Beckman Coulter Genomics). After performing library quality control using the 2100 Bioanalyzer with the DNA High Sensitivity kit (Agilent Technologies), the cDNA was pooled, purified and sequenced using an Illumina NextSeq 500 system with 10 million reads/sample in single end mode with 75 nt read length. RNA-seq raw reads were trimmed, filtered and mapped using the *F. nucleatum* ssp. *nucleatum* ATCC 23726 (NZ_CP028109.1) annotation file with custom annotation and for *F. nucleatum* ssp. *vincentii* ATCC 49256 the annotation file of *F. nucleatum* ssp. *vincentii* 3_1_36A2 (NZ_ CP003700.1). To remove adapter sequences and trim nucleobases BBDuk was used before mapping the reads with BBMap (v38.84). Differential gene expression was investigated using the featureCounts methods of the Subread (2.0.1) package. For further downstream analysis R/Bioconductor packages were used. The reads were normalized by the trimmed mean of *M* values (TMM) normalization. Data of three biological replicates showing a log_2_ fold change <−1.8 or >1.8 with a false-discovery-rate (FDR) ≤ 0.01 were considered as differentially regulated transcripts.

### KEGG pathway enrichment analysis

Genes were assigned to KEGG pathways using the R package KEGGREST (1.32.0). To investigate enrichment of pathways in differentially expressed transcripts, rotation gene set testing was applied. All pathways with more than ten transcripts and an FDR-adjusted P-value <0.001 were visualized as statistically significant via asterisk. The color in the heat map indicates the median log_2_ FC of the respective pathway.

### Data Availability

RNA-seq data can be accessed at NCBI’s GEO (https://www.ncbi.nlm.nih.gov/geo) under the accession number GSE284320.

### Code Availability

The code used for RNA-seq data analysis is available at https://github.com/jakobjung/fuso_PNA_rnaseq.

### Author Contributions

V.C. performed all of the experiments. J.J. performed data analysis of RNA-seq. C.G. coupled PMOs to CPPs and performed HPLC analysis of the compounds. V.C., L.P., and F.P. designed research. J.V. directed research. V.C., J.J., F.P., L.P., L.B., and J.V. wrote the manuscript.

## Acknowledgements

We are grateful to Anke Sparmann for her great help on editing of the manuscript. Additionally, we would like to thank Anna Nöhren and Esther Hauschild for their stellar technical assistance. Further, we are grateful to Mark Brönstrup, Werner Tegge, and Brigitte Kornak for synthetizing CPP-PNAs at the HZI. We thank the Vogel Stiftung Dr. Eckernkamp for supporting V.C. and F.P. with a Dr. Eckernkamp Fellowship. This work was supported by funds to J.V. from a DFG Gottfried Wilhelm Leibniz Award (DFG Vo875-18) and the Bavarian bayresq.net project Rbiotics (L.B., J.V.). Research was further funded by the BMBF in the framework of the Cluster4Future program (Cluster for Nucleic Acid Therapeutics Munich, CNATM; Project ID: 03ZU1201CA; L.P., J.V.) and the Deutsche Forschungsgemeinschaft (DFG; SFB 1583/1, Project number: 492620490, Subproject A09; J.V.).

## Supplementary figures

**Fig. S1:**
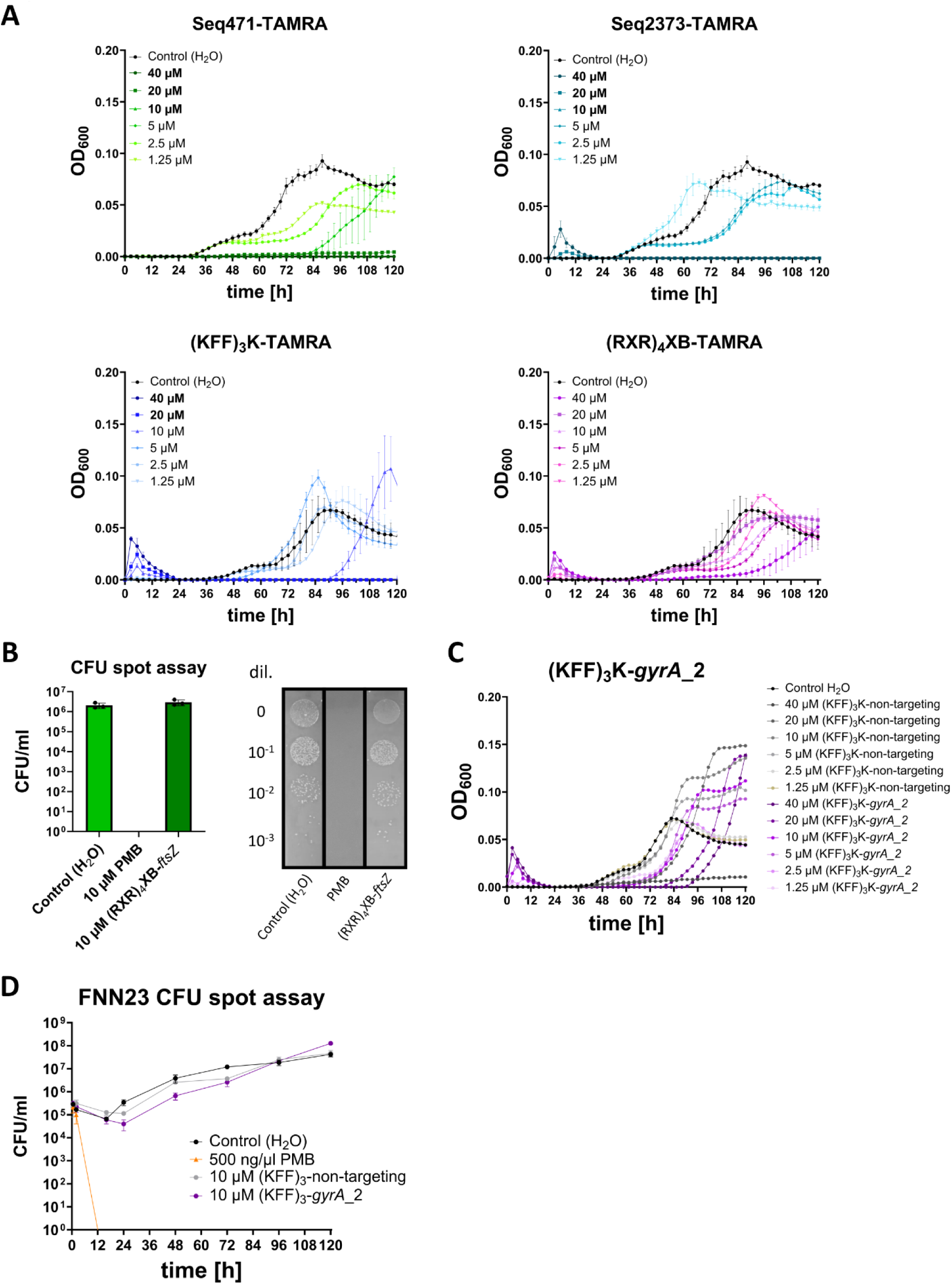
MIC determination of CPP-TAMRA conjugates and investigation of (RXR)_4_XB-*ftsZ* as well as (KFF)_3_K-*gyrA*_2. (A) Growth kinetics of FNN23 incubated with CPP-TAMRA conjugates Seq471, Seq2373, (KFF)_3_K or (RXR)_4_XB in MHB with 1 x 10^5^ CFU per ml input. Growth curves are depicted as OD_600nm_ over time for three technical replicates, error bars indicate the standard deviation. (B) Spot assay of FNN23 incubated with 10 µM (RXR)_4_XB-*ftsZ* to test its effect on colony formation together with H_2_O as negative control and PMB as positive control. At 96 h post treatment, 100 µl of each sample were collected and diluted 1:10 in 1x PBS to enumerate the number of viable cells (CFU/ml) via spotting on BHI agar plates. (Left) After three days CFUs were counted for three technical replicates and quantified. Error bars represent standard deviation of three experiments. (Right) Exemplary images out of three replicates showing CFUs on BHI plates from treated samples. (C) Growth kinetics of FNN23 incubated with water, (KFF)_3_K-*gyrA*_2 or (KFF)_3_K-non-targeting control in MHB with 1 x 10^5^ CFU per ml input. Growth curves are depicted as OD_600nm_ over time. (D) Determination of bactericidal kinetics for 10 µM (KFF)_3_K-*gyrA*_2 or (KFF)_3_K-non-targeting control as well as H_2_O as negative control and 10 µM PMB as positive control. At selected time points post treatment, 100 µl of each sample were collected and diluted 1:10 in 1x PBS to enumerate the number of viable cells (CFU/ml) via spotting on BHI agar plates. After three days incubation at 37°C CFUs on plates were counted and quantified for each condition. Error bars represent standard deviation of three experiments.

**Fig. S2:**
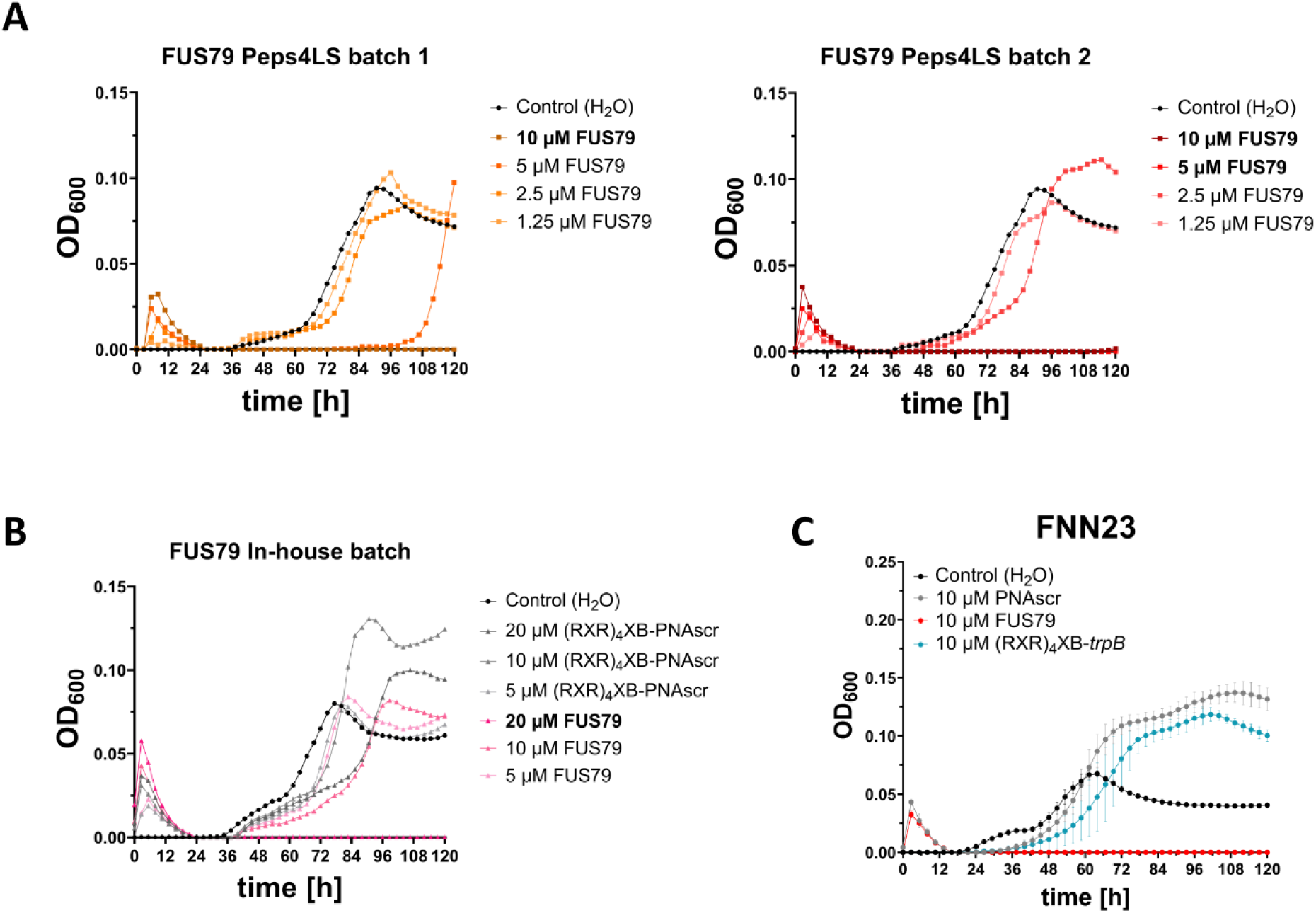
Comparison of different batches FUS79 and *trpB* off-target investigation. (A) Growth kinetics of FNN23 incubated with different Peps4LS batches of FUS79 or water as control. (B) Growth kinetics of FNN23 incubated with in house synthetized FUS79 or corresponding scrambled control (PNAscr), or water as control. (C) Growth kinetics of FNN23 incubated with 10 µM FUS79 (red), PNAscr (gray) or (RXR)_4_XB-PNA complementary to TIR off-target *trpB* (green). Error bars represent the standard deviation of three experiments. All growth curves are depicted as OD_600nm_ over time with 1 x 10^5^ CFU per ml input in MHB.

**Fig S3:**
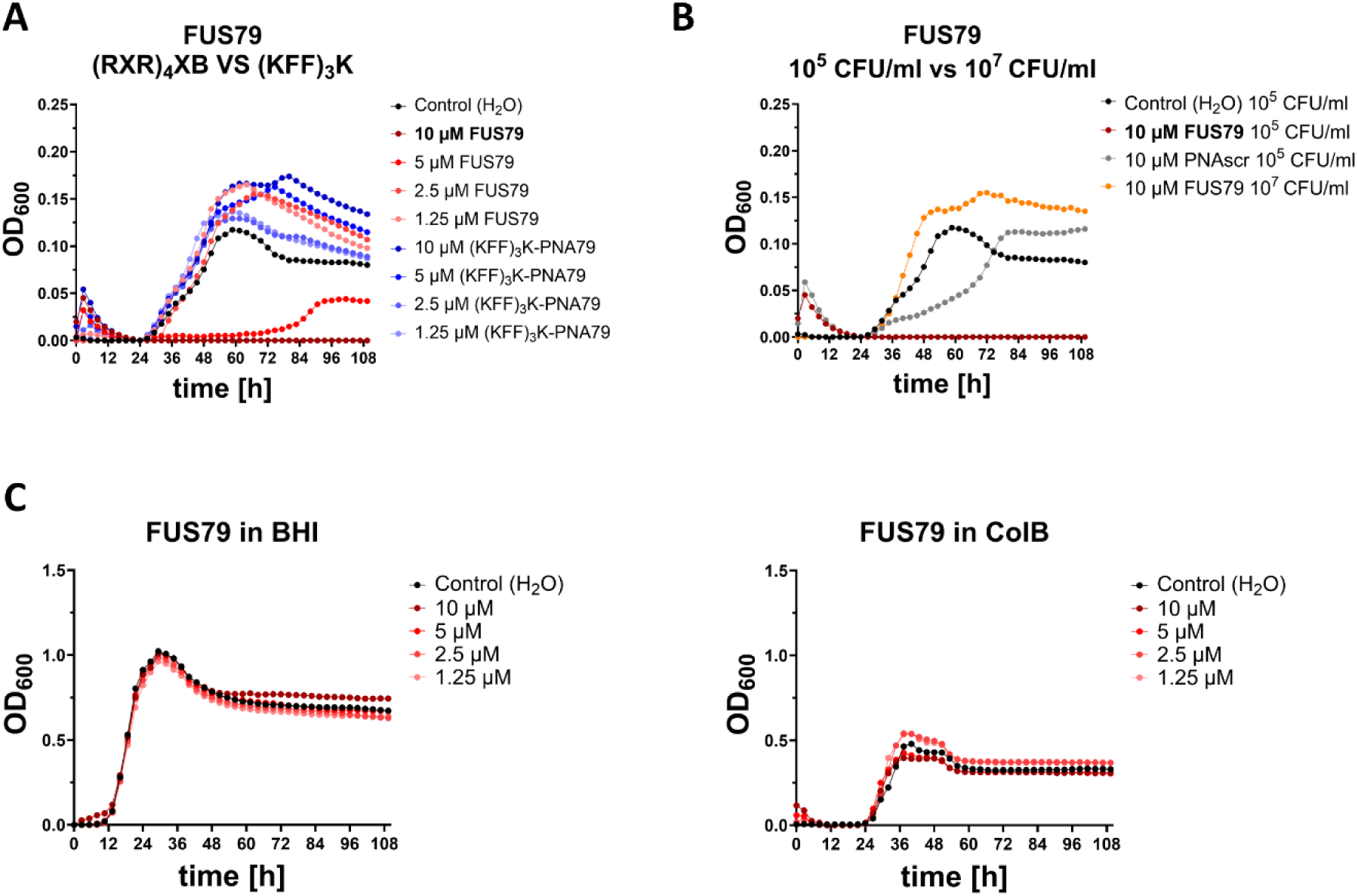
Investigation of CPP-, inoculum- and media-dependency of the antibacterial activity of FUS79. (A) Growth kinetics of FNN23 incubated with FUS79 (=(RXR)_4_XB-PNA79, red), (KFF)_3_K-PNA79 (blue), or water control (black) in MHB with 1 x 10^5^ CFU per ml input. (B) Growth kinetics of FNN23 in MHB incubated with 10 µM FUS79 (red), PNAscr (gray) or water control (black) using 1 x 10^5^ CFU versus 10 µM FUS79 using 1 x 10^7^ CFU per ml input (orange). (C) Growth kinetics of FNN23 incubated with FUS79 (red) or water (black) in cation- and peptide-rich growth media using 1 x 10^5^ CFU per ml input: BHI (left) and ColB (right). All growth curves depicted as OD_600nm_ over time.

**Fig. S4:**
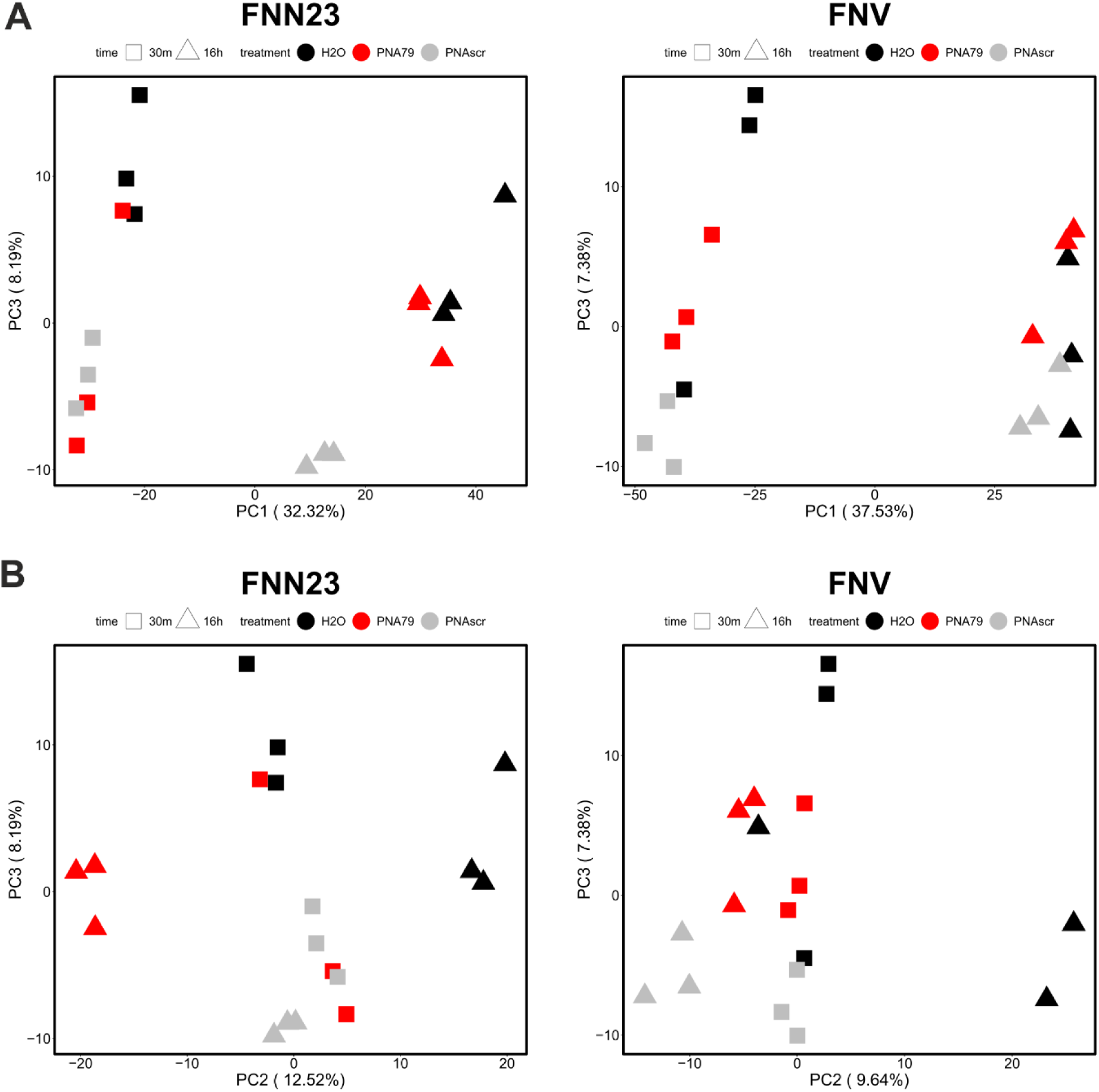
Principal component analysis of RNA-seq: PC1 vs PC3 as well as PC2 vs PC3. Principal component analysis (PCA) of 18 samples for each fusobacterial strain after trimmed mean of M-values (TMM) normalization at 30 minutes and 16 hours after treatment. (A) PC1 vs PC3 and (B) PC2 vs PC3.

**Fig. S5:**
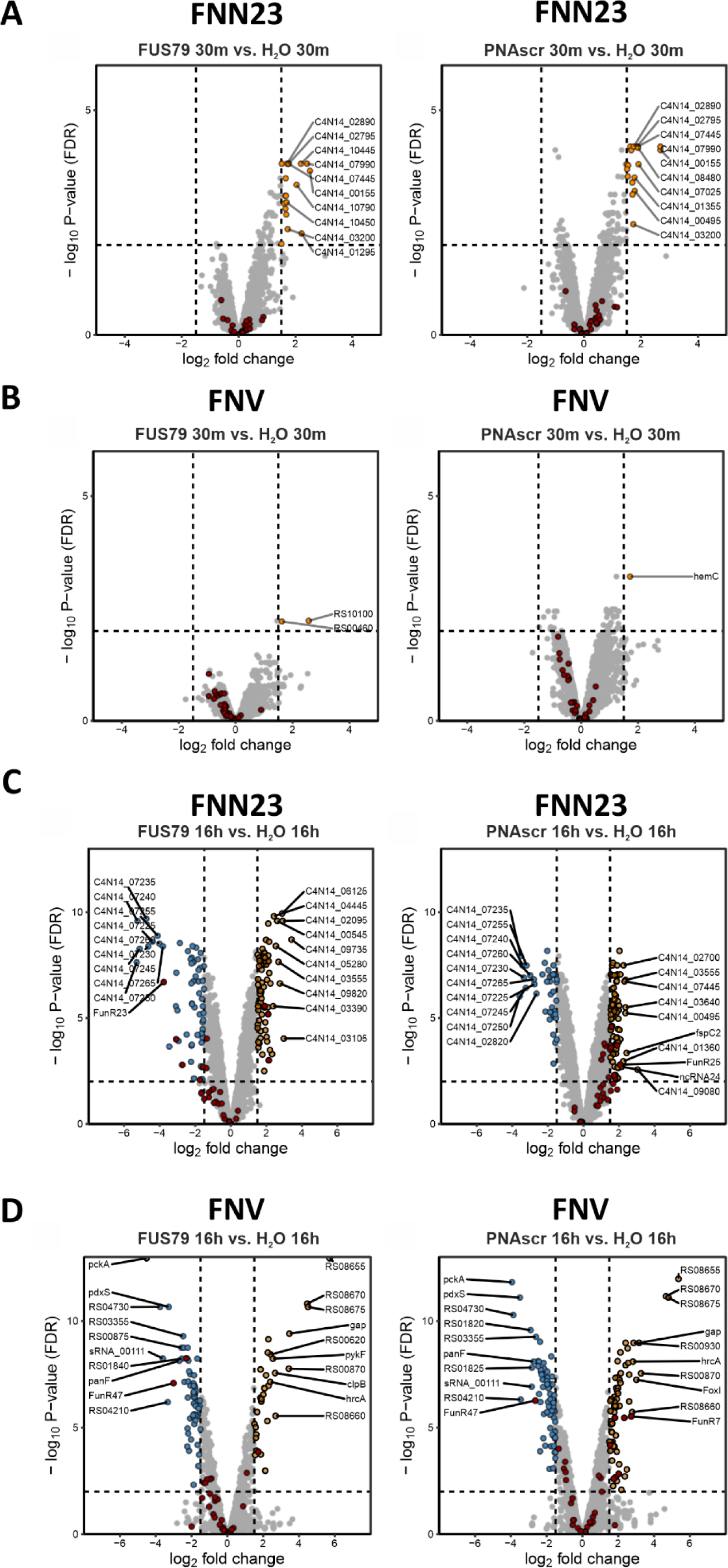
Transcriptomic profiling of FNN23 and FNV in response to (RXR)_4_XB-PNA treatment. Transcriptomic response of FNN23 (A) and FNV (B) upon PNA-FUS79 or respective PNAscr treatment compared to H_2_O control at 30 minutes and transcriptomic response of FNN23 (C) and FNV (D) upon PNA-FUS79 or PNAscr treatment compared to H_2_O control at 16 hours. For A-D volcano plots show differential gene expression as –log_10_ false discovery rate (FDR)-adjusted P-values on y-axis and log_2_ fold change on x-axis. Significantly differentially expressed transcripts are defined by an absolute fold change <-1.8 or >1.8 and an FDR adjusted P-value <0.01, characterized by the dashed lines in the plot. Significantly upregulated transcripts are depicted in orange, significantly downregulated transcripts in blue and all sRNAs dots are colored in red. The top ten differentially expressed transcripts are specified by the depicted locus tag.

**Fig. S6:**
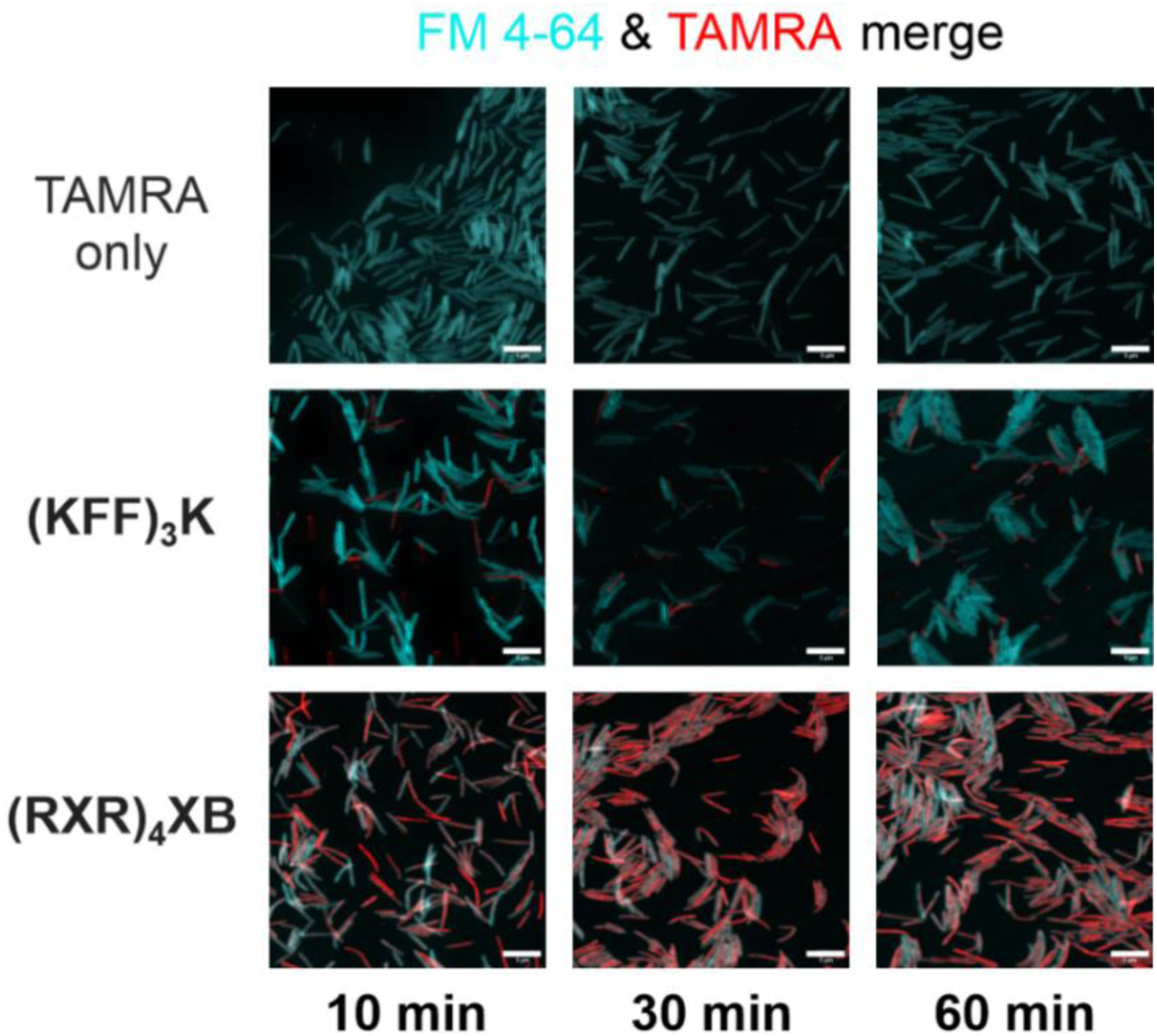
Investigation of CPP-TAMRA penetration in FNV using CLSM. Representative CLSM images for TAMRA only control or TAMRA-labeled CPPs (red) at 5 µM and 10, 30 and 60 minutes post incubation. FNV was incubated with FM 4-64 to stain the cell membrane, shown in cyan. Scale bar, 5 µm.

## Supplementary Tables

**Table S1:**
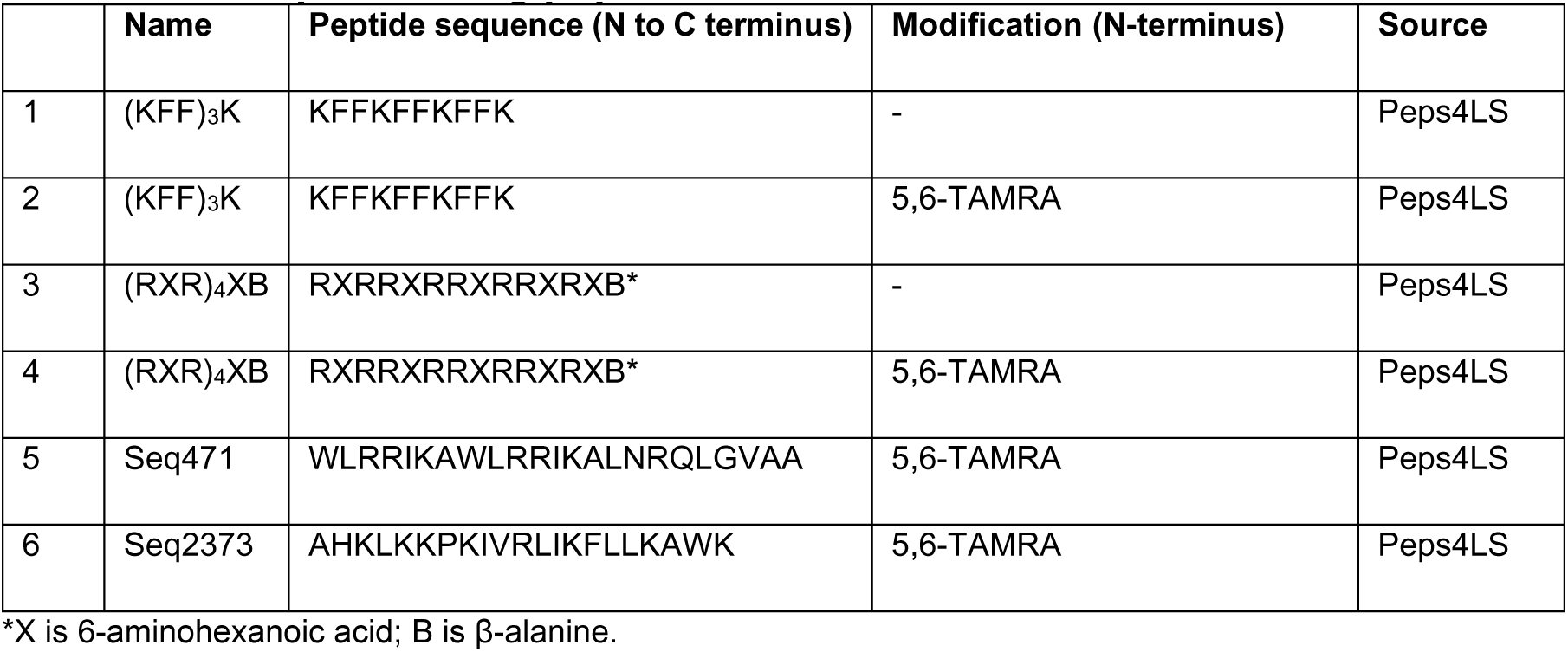
Cell penetrating peptides.

**Table S2:**
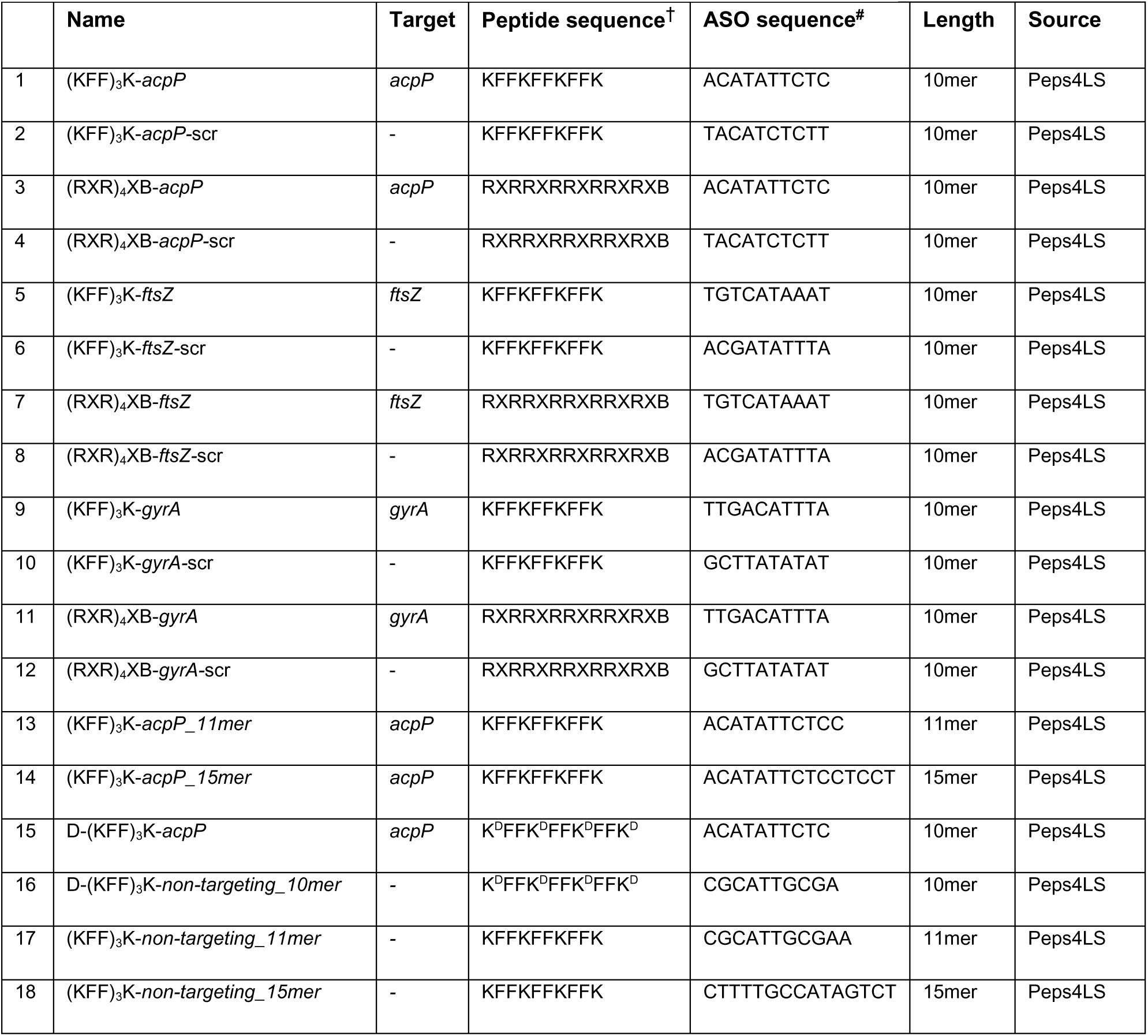
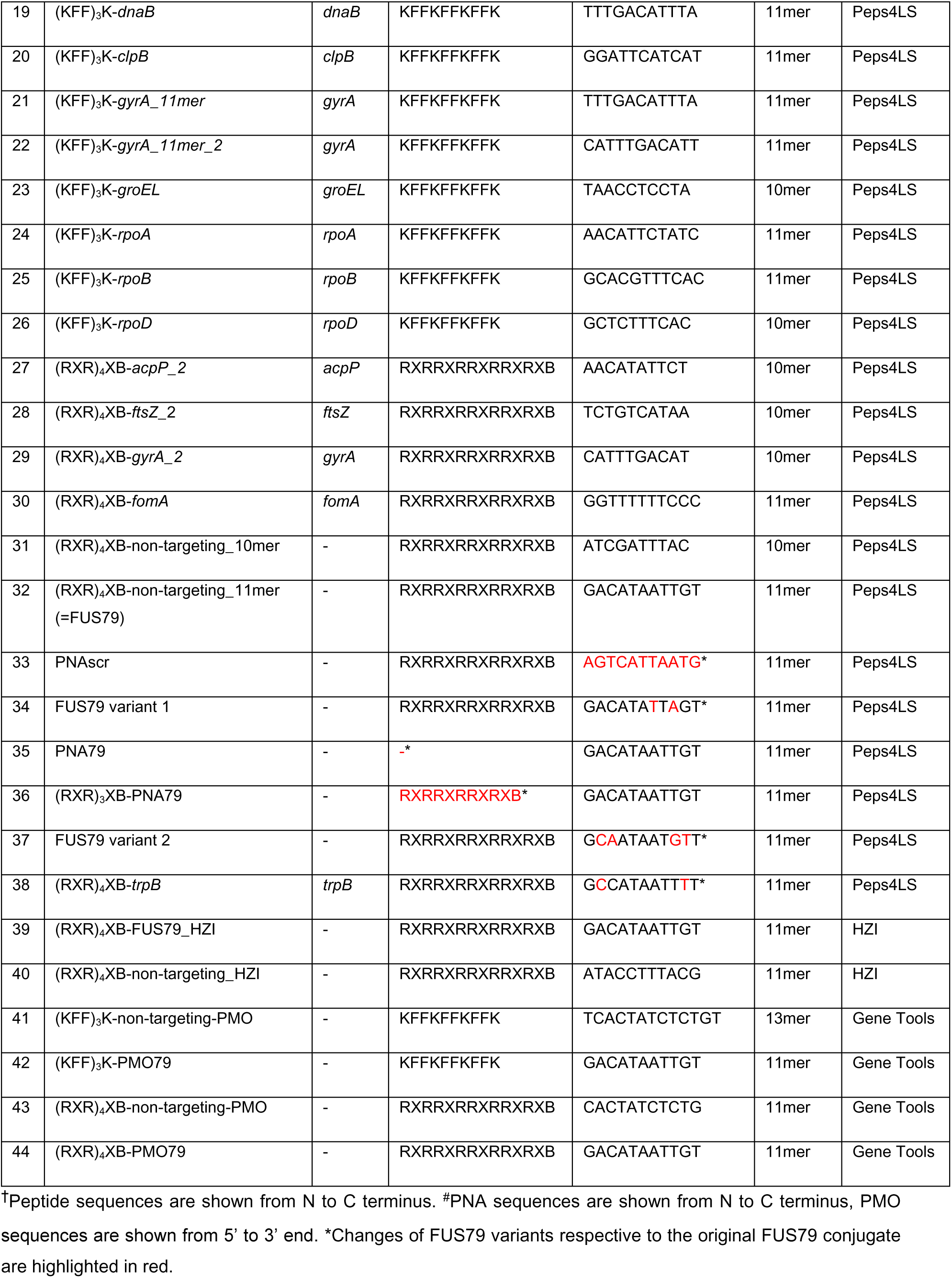
Antisense oligomer conjugates.

## REFERENCES

1. Baker JL, Mark Welch JL, Kauffman KM, McLean JS, He X. 2024. The oral microbiome: diversity, biogeography and human health. Nat Rev Microbiol 22:89–104.

2. Segata N, Haake SK, Mannon P, Lemon KP, Waldron L, Gevers D, Huttenhower C, Izard J. 2012. Composition of the adult digestive tract bacterial microbiome based on seven mouth surfaces, tonsils, throat and stool samples. Genome Biol 13:R42.

3. Kaplan CW, Ma X, Paranjpe A, Jewett A, Lux R, Kinder-Haake S, Shi W. 2010. Fusobacterium nucleatum Outer Membrane Proteins Fap2 and RadD Induce Cell Death in Human Lymphocytes. Infect Immun 78:4773–4778.

4. Nithianantham S, Xu M, Yamada M, Ikegami A, Shoham M, Han YW. 2009. Crystal Structure of FadA Adhesin from Fusobacterium nucleatum Reveals a Novel Oligomerization Motif, the Leucine Chain. J Biol Chem 284:3865–3872.

5. Nakagaki H, Sekine S, Terao Y, Toe M, Tanaka M, Ito H-O, Kawabata S, Shizukuishi S, Fujihashi K, Kataoka K. 2010. Fusobacterium nucleatum Envelope Protein FomA Is Immunogenic and Binds to the Salivary Statherin-Derived Peptide. Infect Immun 78:1185–1192.

6. Brennan CA, Garrett WS. 2019. Fusobacterium nucleatum — symbiont, opportunist and oncobacterium. Nat Rev Microbiol 17:156–166.

7. Kaplan CW, Lux R, Haake SK, Shi W. 2009. The Fusobacterium nucleatum outer membrane protein RadD is an arginine-inhibitable adhesin required for inter-species adherence and the structured architecture of multispecies biofilm. Mol Microbiol 71:35–47.

8. Mark Welch JL, Rossetti BJ, Rieken CW, Dewhirst FE, Borisy GG. 2016. Biogeography of a human oral microbiome at the micron scale. Proc Natl Acad Sci U S A 113:E791–800.

9. Han YW. 2015. Fusobacterium nucleatum: a commensal-turned pathogen. Curr Opin Microbiol 0:141–147.

10. Pignatelli P, Nuccio F, Piattelli A, Curia MC. 2023. The Role of Fusobacterium nucleatum in Oral and Colorectal Carcinogenesis. Microorganisms 11:2358.

11. Abed J, Maalouf N, Manson AL, Earl AM, Parhi L, Emgård JEM, Klutstein M, Tayeb S, Almogy G, Atlan KA, Chaushu S, Israeli E, Mandelboim O, Garrett WS, Bachrach G. 2020. Colon Cancer-Associated Fusobacterium nucleatum May Originate From the Oral Cavity and Reach Colon Tumors via the Circulatory System. Front Cell Infect Microbiol 10:400.

12. Kostic AD, Chun E, Robertson L, Glickman JN, Gallini CA, Michaud M, Clancy TE, Chung DC, Lochhead P, Hold GL, El-Omar EM, Brenner D, Fuchs CS, Meyerson M, Garrett WS. 2013. *Fusobacterium nucleatum* Potentiates Intestinal Tumorigenesis and Modulates the Tumor-Immune Microenvironment. Cell Host Microbe 14:207–215.

13. Ou S, Chen H, Wang H, Ye J, Liu H, Tao Y, Ran S, Mu X, Liu F, Zhu S, Luo K, Guan Z, Jin Y, Huang R, Song Y, Liu S. 2023. Fusobacterium nucleatum upregulates MMP7 to promote metastasis-related characteristics of colorectal cancer cell via activating MAPK(JNK)-AP1 axis. J Transl Med 21:704.

14. Yu T, Guo F, Yu Y, Sun T, Ma D, Han J, Qian Y, Kryczek I, Sun D, Nagarsheth N, Chen Y, Chen H, Hong J, Zou W, Fang J-Y. 2017. *Fusobacterium nucleatum* Promotes Chemoresistance to Colorectal Cancer by Modulating Autophagy. Cell 170:548–563.e16.

15. Bullman S, Pedamallu CS, Sicinska E, Clancy TE, Zhang X, Cai D, Neuberg D, Huang K, Guevara F, Nelson T, Chipashvili O, Hagan T, Walker M, Ramachandran A, Diosdado B, Serna G, Mulet N, Landolfi S, Cajal SR y, Fasani R, Aguirre AJ, Ng K, Élez E, Ogino S, Tabernero J, Fuchs CS, Hahn WC, Nuciforo P, Meyerson M. 2017. Analysis of Fusobacterium persistence and antibiotic response in colorectal cancer. Science 358:1443–1448.

16. Parhi L, Alon-Maimon T, Sol A, Nejman D, Shhadeh A, Fainsod-Levi T, Yajuk O, Isaacson B, Abed J, Maalouf N, Nissan A, Sandbank J, Yehuda-Shnaidman E, Ponath F, Vogel J, Mandelboim O, Granot Z, Straussman R, Bachrach G. 2020. Breast cancer colonization by Fusobacterium nucleatum accelerates tumor growth and metastatic progression. Nat Commun 11:3259.

17. Gao Y, Shang Q, Li W, Guo W, Stojadinovic A, Mannion C, Man Y, Chen T. 2020. Antibiotics for cancer treatment: A double-edged sword. J Cancer 11:5135–5149.

18. Angrish N, Khare G. 2023. Antisense oligonucleotide based therapeutics and its applications against bacterial infections. Med Drug Discov 20:100166.

19. Moreira L, Guimarães NM, Santos RS, Loureiro JA, Pereira MC, Azevedo NF. 2024. Promising strategies employing nucleic acids as antimicrobial drugs. Mol Ther - Nucleic Acids 35.

20. Vogel J. 2020. An RNA biology perspective on species-specific programmable RNA antibiotics. Mol Microbiol 113:550–559.

21. Vogel J, Faber F, Barquist L, Sparmann A, Popella L, Ghosh C. In press. Meeting report ASOBIOTICS 2024: an interdisciplinary symposium on antisense-based programmable RNA antibiotics. RNA.

22. Pradeep SP, Malik S, Slack FJ, Bahal R. 2023. Unlocking the potential of chemically modified peptide nucleic acids for RNA-based therapeutics. RNA 29:434–445.

23. Nielsen PE, Egholm M, Berg RH, Buchardt O. 1991. Sequence-selective recognition of DNA by strand displacement with a thymine-substituted polyamide. Science 254:1497–1500.

24. Ghosh C, Popella L, Dhamodharan V, Jung J, Dietzsch J, Barquist L, Höbartner C, Vogel J. 2024. A comparative analysis of peptide-delivered antisense antibiotics using diverse nucleotide mimics. RNA 30:624–643.

25. Popella L, Jung J, Do PT, Hayward RJ, Barquist L, Vogel J. 2022. Comprehensive analysis of PNA-based antisense antibiotics targeting various essential genes in uropathogenic Escherichia coli. Nucleic Acids Res 50:6435–6452.

26. Good L, Awasthi SK, Dryselius R, Larsson O, Nielsen PE. 2001. Bactericidal antisense effects of peptide-PNA conjugates. Nat Biotechnol 19:360–364.

27. Geller BL, Deere JD, Stein DA, Kroeker AD, Moulton HM, Iversen PL. 2003. Inhibition of gene expression in Escherichia coli by antisense phosphorodiamidate morpholino oligomers. Antimicrob Agents Chemother 47:3233–3239.

28. Nielsen PE, Shiraishi T. 2011. Peptide nucleic acid (PNA) cell penetrating peptide (CPP) conjugates as carriers for cellular delivery of antisense oligomers. Artif DNA PNA XNA 2:90–99.

29. Szabó I, Yousef M, Soltész D, Bató C, Mező G, Bánóczi Z. 2022. Redesigning of Cell-Penetrating Peptides to Improve Their Efficacy as a Drug Delivery System. Pharmaceutics 14:907.

30. Ghosal A, Vitali A, Stach JEM, Nielsen PE. 2013. Role of SbmA in the uptake of peptide nucleic acid (PNA)-peptide conjugates in E. coli. ACS Chem Biol 8:360–367.

31. Barkowsky G, Lemster A-L, Pappesch R, Jacob A, Krüger S, Schröder A, Kreikemeyer B, Patenge N. 2019. Influence of Different Cell-Penetrating Peptides on the Antimicrobial Efficiency of PNAs in Streptococcus pyogenes. Mol Ther Nucleic Acids 18:444–454.

32. Popella L, Jung J, Popova K, Ðurica-Mitić S, Barquist L, Vogel J. 2021. Global RNA profiles show target selectivity and physiological effects of peptide-delivered antisense antibiotics. Nucleic Acids Res 49:4705–4724.

33. El-Fateh M, Chatterjee A, Zhao X. 2024. A systematic review of peptide nucleic acids (PNAs) with antibacterial activities: Efficacy, potential and challenges. Int J Antimicrob Agents 63:107083.

34. Wojciechowska M, Równicki M, Mieczkowski A, Miszkiewicz J, Trylska J. 2020. Antibacterial Peptide Nucleic Acids—Facts and Perspectives. 3. Molecules 25:559.

35. Sugimoto S, Maeda H, Kitamatsu M, Nishikawa I, Shida M. 2019. Selective growth inhibition of Porphyromonas gingivalis and Aggregatibacter actinomycetemcomitans by antisense peptide nucleic acids. Mol Cell Probes 43:45–49.

36. Lee H-M, Ren J, Tran KM, Jeon B-M, Park W-U, Kim H, Lee KE, Oh Y, Choi M, Kim D-S, Na D. 2021. Identification of efficient prokaryotic cell-penetrating peptides with applications in bacterial biotechnology. Commun Biol 4:205.

37. Ramaker K, Henkel M, Krause T, Röckendorf N, Frey A. 2018. Cell penetrating peptides: a comparative transport analysis for 474 sequence motifs. Drug Deliv 25:928–937.

38. Wojciechowska M, Miszkiewicz J, Trylska J. 2020. Conformational Changes of Anoplin, W-MreB1–9, and (KFF)3K Peptides near the Membranes. Int J Mol Sci 21:9672.

39. Drexelius M, Reinhardt A, Grabeck J, Cronenberg T, Nitsche F, Huesgen PF, Maier B, Neundorf I. 2021. Multistep optimization of a cell-penetrating peptide towards its antimicrobial activity. Biochem J 478:63–78.

40. Birch D, Christensen MV, Staerk D, Franzyk H, Nielsen HM. 2017. Fluorophore labeling of a cell-penetrating peptide induces differential effects on its cellular distribution and affects cell viability. Biochim Biophys Acta BBA - Biomembr 1859:2483–2494.

41. Good L, Nielsen PE. 1998. Antisense inhibition of gene expression in bacteria by PNA targeted to mRNA. Nat Biotechnol 16:355–358.

42. A. Ghomi F, Jung JJ, Langridge GC, Cain AK, Boinett CJ, Abd El Ghany M, Pickard DJ, Kingsley RA, Thomson NR, Parkhill J, Gardner PP, Barquist L. 2024. High-throughput transposon mutagenesis in the family Enterobacteriaceae reveals core essential genes and rapid turnover of essentiality. mBio 15:e01798-24.

43. Ponath F, Tawk C, Zhu Y, Barquist L, Faber F, Vogel J. 2021. RNA landscape of the emerging cancer-associated microbe Fusobacterium nucleatum. Nat Microbiol 6:1007–1020.

44. Ponath F, Zhu Y, Vogel J. 2024. Transcriptome fine-mapping in Fusobacterium nucleatum reveals FoxJ, a new σE-dependent small RNA with unusual mRNA activation activity. mBio 15:e0353623.

45. Jung J, Popella L, Do PT, Pfau P, Vogel J, Barquist L. 2023. Design and off-target prediction for antisense oligomers targeting bacterial mRNAs with the MASON web server. RNA 29:570–583.

46. Zhou P, G. C. B, Stolte F, Wu C. 2024. Use of CRISPR interference for efficient and rapid gene inactivation in Fusobacterium nucleatum. Appl Environ Microbiol 90:e01665–23.

47. Goltermann L, Yavari N, Zhang M, Ghosal A, Nielsen PE. 2019. PNA Length Restriction of Antibacterial Activity of Peptide-PNA Conjugates in Escherichia coli Through Effects of the Inner Membrane. Front Microbiol 10:1032.

48. Hadjicharalambous A, Bournakas N, Newman H, Skynner MJ, Beswick P. 2022. Antimicrobial and Cell-Penetrating Peptides: Understanding Penetration for the Design of Novel Conjugate Antibiotics. 11. Antibiotics 11:1636.

49. Yavari N, Goltermann L, Nielsen PE. 2021. Uptake, Stability, and Activity of Antisense Anti-acpP PNA-Peptide Conjugates in Escherichia coli and the Role of SbmA. ACS Chem Biol 16:471–479.

50. Abushahba MFN, Mohammad H, Thangamani S, Hussein AAA, Seleem MN. 2016. Impact of different cell penetrating peptides on the efficacy of antisense therapeutics for targeting intracellular pathogens. Sci Rep 6:20832.

51. Martin-Gallausiaux C, Malabirade A, Habier J, Wilmes P. 2020. Fusobacterium nucleatum Extracellular Vesicles Modulate Gut Epithelial Cell Innate Immunity via FomA and TLR2. Front Immunol 11:583644.

52. Chen Y, Huang Z, Tang Z, Huang Y, Huang M, Liu H, Ziebolz D, Schmalz G, Jia B, Zhao J. 2022. More Than Just a Periodontal Pathogen –the Research Progress on Fusobacterium nucleatum. Front Cell Infect Microbiol 12.

53. Wang B, Deng J, Donati V, Merali N, Frampton AE, Giovannetti E, Deng D. 2024. The Roles and Interactions of Porphyromonas gingivalis and Fusobacterium nucleatum in Oral and Gastrointestinal Carcinogenesis: A Narrative Review. 1. Pathogens 13:93.

54. Martinson JNV, Walk ST. 2020. Escherichia coli Residency in the Gut of Healthy Human Adults. EcoSal Plus 9.

55. Zhong Y, Wilkinson-White L, Zhang E, Mohanty B, Zhang BB, McRae MS, Luo R, Allport TA, Duff AP, Zhao J, El-Kamand S, Du Plessis M-D, Cubeddu L, Gamsjaeger R, Ataide SF, Kwan AH. 2024. Peptide nucleic acids can form hairpins and bind RNA-binding proteins. PloS One 19:e0310565.

56. Loffredo MR, Savini F, Bobone S, Casciaro B, Franzyk H, Mangoni ML, Stella L. 2021. Inoculum effect of antimicrobial peptides. Proc Natl Acad Sci 118:e2014364118.

57. Balouiri M, Sadiki M, Ibnsouda SK. 2016. Methods for in vitro evaluating antimicrobial activity: A review. J Pharm Anal 6:71–79.

58. Hudziak RM, Summerton J, Weller DD, Iversen PL. 2000. Antiproliferative effects of steric blocking phosphorodiamidate morpholino antisense agents directed against c-myc. Antisense Nucleic Acid Drug Dev 10:163–176.

59. Sun X, Marque LO, Cordner Z, Pruitt JL, Bhat M, Li PP, Kannan G, Ladenheim EE, Moran TH, Margolis RL, Rudnicki DD. 2014. Phosphorodiamidate morpholino oligomers suppress mutant huntingtin expression and attenuate neurotoxicity. Hum Mol Genet 23:6302–6317.

60. Ponath F, Zhu Y, Cosi V, Vogel J. 2022. Expanding the genetic toolkit helps dissect a global stress response in the early-branching species Fusobacterium nucleatum. Proc Natl Acad Sci 119:e2201460119.

61. Rowley G, Spector M, Kormanec J, Roberts M. 2006. Pushing the envelope: extracytoplasmic stress responses in bacterial pathogens. Nat Rev Microbiol 4:383– 394.

62. Hayden JD, Ades SE. 2008. The Extracytoplasmic Stress Factor, σE, Is Required to Maintain Cell Envelope Integrity in Escherichia coli. PLOS ONE 3:e1573.

63. Brown S, Fournier MJ. 1984. The 4.5 S RNA gene of Escherichia coli is essential for cell growth. J Mol Biol 178:533–550.

64. Yang AJ, Mulligan RM. 1996. Identification of a 4.5S-like ribonucleoprotein in maize mitochondria. Nucleic Acids Res 24:3601–3606.

65. Xiong L, Teng JLL, Botelho MG, Lo RC, Lau SKP, Woo PCY. 2016. Arginine Metabolism in Bacterial Pathogenesis and Cancer Therapy. Int J Mol Sci 17.

66. Christgen SL, Becker DF. 2019. Role of Proline in Pathogen and Host Interactions. Antioxid Redox Signal 30:683–709.

67. Bie L, Zhang M, Wang J, Fang M, Li L, Xu H, Wang M. 2023. Comparative Analysis of Transcriptomic Response of Escherichia coli K-12 MG1655 to Nine Representative Classes of Antibiotics. Microbiol Spectr 11:e0031723.

68. Galinier A, Deutscher J. 2017. Sophisticated Regulation of Transcriptional Factors by the Bacterial Phosphoenolpyruvate: Sugar Phosphotransferase System. J Mol Biol 429:773–789.

69. Masip L, Veeravalli K, Georgiou G. 2006. The many faces of glutathione in bacteria. Antioxid Redox Signal 8:753–762.

70. Hartl J, Kiefer P, Kaczmarczyk A, Mittelviefhaus M, Meyer F, Vonderach T, Hattendorf B, Jenal U, Vorholt JA. 2020. Untargeted metabolomics links glutathione to bacterial cell cycle progression. Nat Metab 2:153–166.

71. Zhu Y, Ponath F, Cosi V, Vogel J. 2024. A global survey of small RNA interactors identifies KhpA and KhpB as major RNA-binding proteins in Fusobacterium nucleatum. Nucleic Acids Res 52:3950–3970.

72. Desnoyers G, Bouchard M-P, Massé E. 2013. New insights into small RNA-dependent translational regulation in prokaryotes. Trends Genet 29:92–98.

73. Fröhlich KS, Papenfort K. 2020. Regulation outside the box: New mechanisms for small RNAs. Mol Microbiol 114:363–366.

74. Bai H, You Y, Yan H, Meng J, Xue X, Hou Z, Zhou Y, Ma X, Sang G, Luo X. 2012. Antisense inhibition of gene expression and growth in gram-negative bacteria by cell-penetrating peptide conjugates of peptide nucleic acids targeted to *rpoD* gene. Biomaterials 33:659–667.

75. Martin I, Underhaug J, Celaya G, Moro F, Teigen K, Martinez A, Muga A. 2013. Screening and Evaluation of Small Organic Molecules as ClpB Inhibitors and Potential Antimicrobials. J Med Chem 56:7177–7189.

76. Soofi MA, Seleem MN. 2012. Targeting Essential Genes in Salmonella enterica Serovar Typhimurium with Antisense Peptide Nucleic Acid. Antimicrob Agents Chemother 56:6407–6409.

77. Abushahba MF, Mohammad H, Seleem MN. 2016. Targeting Multidrug-resistant Staphylococci with an anti-rpoA Peptide Nucleic Acid Conjugated to the HIV-1 TAT Cell Penetrating Peptide. Mol Ther Nucleic Acids 5:e339.

78. Tan KXY, Shigenobu S. 2024. In vivo interference of pea aphid endosymbiont Buchnera groEL gene by synthetic peptide nucleic acids. Sci Rep 14:5378.

79. Wu C, Chen Y-W, Scheible M, Chang C, Wittchen M, Lee JH, Luong TT, Tiner BL, Tauch A, Das A, Ton-That H. 2021. Genetic and molecular determinants of polymicrobial interactions in Fusobacterium nucleatum. Proc Natl Acad Sci 118:e2006482118.

80. Kumar A, Thotakura PL, Tiwary BK, Krishna R. 2016. Target identification in Fusobacterium nucleatum by subtractive genomics approach and enrichment analysis of host-pathogen protein-protein interactions. BMC Microbiol 16:84.

81. Cain AK, Barquist L, Goodman AL, Paulsen IT, Parkhill J, van Opijnen T. 2020. A decade of advances in transposon-insertion sequencing. Nat Rev Genet 21:526–540.

82. Coppenhagen-Glazer S, Sol A, Abed J, Naor R, Zhang X, Han YW, Bachrach G. 2015. Fap2 of Fusobacterium nucleatum Is a Galactose-Inhibitable Adhesin Involved in Coaggregation, Cell Adhesion, and Preterm Birth. Infect Immun 83:1104–1113.

83. Kapatral V, Anderson I, Ivanova N, Reznik G, Los T, Lykidis A, Bhattacharyya A, Bartman A, Gardner W, Grechkin G, Zhu L, Vasieva O, Chu L, Kogan Y, Chaga O, Goltsman E, Bernal A, Larsen N, D’Souza M, Walunas T, Pusch G, Haselkorn R, Fonstein M, Kyrpides N, Overbeek R. 2002. Genome Sequence and Analysis of the Oral Bacterium Fusobacterium nucleatum Strain ATCC 25586. J Bacteriol 184:2005– 2018.

84. Zhang Y, Xie X, Ma W, Zhan Y, Mao C, Shao X, Lin Y. 2020. Multi-targeted Antisense Oligonucleotide Delivery by a Framework Nucleic Acid for Inhibiting Biofilm Formation and Virulence. Nano-Micro Lett 12:74.

85. Good L. 2002. Antisense inhibition of bacterial gene expression and cell growth. Methods Mol Biol Clifton NJ 208:237–248.

86. Oh E, Zhang Q, Jeon B. 2014. Target optimization for peptide nucleic acid (PNA)-mediated antisense inhibition of the CmeABC multidrug efflux pump in Campylobacter jejuni. J Antimicrob Chemother 69:375–380.

87. Porosk L, Langel Ü. 2022. Approaches for evaluation of novel CPP-based cargo delivery systems. Front Pharmacol 13:1056467.

88. Beha MJ, Ryu JS, Kim YS, Chung HJ. 2021. Delivery of antisense oligonucleotides using multi-layer coated gold nanoparticles to methicillin-resistant *S. aureus* for combinatorial treatment. Mater Sci Eng C 126:112167.

89. Ghaffari E, Rezatofighi SE, Ardakani MR, Rastegarzadeh S. 2019. Delivery of antisense peptide nucleic acid by gold nanoparticles for the inhibition of virus replication. Nanomed 14:1827–1840.

90. Równicki M, Wojciechowska M, Wierzba AJ, Czarnecki J, Bartosik D, Gryko D, Trylska J. 2017. Vitamin B12 as a carrier of peptide nucleic acid (PNA) into bacterial cells. Sci Rep 7:7644.

91. Khasheii B, Mahmoodi P, Mohammadzadeh A. 2021. Siderophores: Importance in bacterial pathogenesis and applications in medicine and industry. Microbiol Res 250:126790.

92. Wang C, Yang D, Wang Y, Ni W. 2022. Cefiderocol for the Treatment of Multidrug-Resistant Gram-Negative Bacteria: A Systematic Review of Currently Available Evidence. Front Pharmacol 13:896971.

93. Pals MJ, Wijnberg L, Yildiz Ç, Velema WA. 2024. Catechol-Siderophore Mimics Convey Nucleic Acid Therapeutics into Bacteria. Angew Chem 136:e202402405.

94. Tsylents U, Burmistrz M, Wojciechowska M, Stępień J, Maj P, Trylska J. 2024. Iron uptake pathway of Escherichia coli as an entry route for peptide nucleic acids conjugated with a siderophore mimic. Front Microbiol 15:1331021.

95. Hastie JL, Carmichael HL, Werner BM, Dunbar KE, Carlson PE. 2023. Clostridioides difficile utilizes siderophores as an iron source and FhuDBGC contributes to ferrichrome uptake. J Bacteriol 205:e00324–23.

96. Li W, Ying X, Lu Q, Chen L. 2012. Predicting sRNAs and Their Targets in Bacteria. Genomics Proteomics Bioinformatics 10:276–284.

97. Kapatral V, Ivanova N, Anderson I, Reznik G, Bhattacharyya A, Gardner WL, Mikhailova N, Lapidus A, Larsen N, D’Souza M, Walunas T, Haselkorn R, Overbeek R, Kyrpides N. 2003. Genome Analysis of F. nucleatum sub spp vincentii and Its Comparison With the Genome of F. nucleatum ATCC 25586. Genome Res 13:1180– 1189.

98. Goltermann L, Zhang M, Ebbensgaard AE, Fiodorovaite M, Yavari N, Løbner-Olesen A, Nielsen PE. 2022. Effects of LPS Composition in Escherichia coli on Antibacterial Activity and Bacterial Uptake of Antisense Peptide-PNA Conjugates. Front Microbiol 13:877377.

99. Vinogradov E, St Michael F, Cox AD. 2018. Structure of the LPS O-chain from Fusobacterium nucleatum strain ATCC 23726 containing a novel 5,7-diamino-3,5,7,9-tetradeoxy-l-gluco-non-2-ulosonic acid presumably having the d-glycero-l-gluco configuration. Carbohydr Res 468:69–72.

100. Alba BM, Gross CA. 2004. Regulation of the Escherichia coli sigma-dependent envelope stress response. Mol Microbiol 52:613–619.

101. Anthony JR, Warczak KL, Donohue TJ. 2005. A transcriptional response to singlet oxygen, a toxic byproduct of photosynthesis. Proc Natl Acad Sci U S A 102:6502– 6507.

102. Ho TD, Ellermeier CD. 2019. Activation of the extracytoplasmic function σ factor σV by lysozyme. Mol Microbiol 112:410–419.

103. Poirel L, Jayol A, Nordmann P. 2017. Polymyxins: Antibacterial Activity, Susceptibility Testing, and Resistance Mechanisms Encoded by Plasmids or Chromosomes. Clin Microbiol Rev 30:557–596.

104. Kumar M, Srivastava S. 2011. Effect of calcium and magnesium on the antimicrobial action of enterocin LR/6 produced by *Enterococcus faecium* LR/6. Int J Antimicrob Agents 37:572–575.

105. Kalafatovic D, Giralt E. 2017. Cell-Penetrating Peptides: Design Strategies beyond Primary Structure and Amphipathicity. Mol J Synth Chem Nat Prod Chem 22:1929.

106. Choi H, Chakraborty S, Liu R, Gellman SH, Weisshaar JC. 2014. Medium Effects on Minimum Inhibitory Concentrations of Nylon-3 Polymers against E. coli. PLoS ONE 9:e104500.

107. Aartsma-Rus A, Vliet L van, Hirschi M, Janson AA, Heemskerk H, Winter CL de, Kimpe S de, Deutekom JC van, Hoen PA ’t, Ommen G-JB van. 2009. Guidelines for Antisense Oligonucleotide Design and Insight Into Splice-modulating Mechanisms. Mol Ther 17:548–553.

108. Patel SG, Sayers EJ, He L, Narayan R, Williams TL, Mills EM, Allemann RK, Luk LYP, Jones AT, Tsai Y-H. 2019. Cell-penetrating peptide sequence and modification dependent uptake and subcellular distribution of green florescent protein in different cell lines. Sci Rep 9:6298.

109. Summerton JE, Weller DD. February 1993. Uncharged morpolino-based polymers having phosphorous containing chiral intersubunit linkages. US5185444A.

110. Chen W, Dong B, Liu W, Liu Z. 2021. Recent Advances in Peptide Nucleic Acids as Antibacterial Agents. Curr Med Chem 28:1104–1125.

111. Kabwe M, Brown TL, Dashper S, Speirs L, Ku H, Petrovski S, Chan HT, Lock P, Tucci J. 2019. Genomic, morphological and functional characterisation of novel bacteriophage FNU1 capable of disrupting Fusobacterium nucleatum biofilms. Sci Rep 9:9107.

112. Yakar N, Unlu O, Cen L, Hasturk H, Chen T, Shi W, He X, Kantarci A. 2024. Targeted elimination of Fusobacterium nucleatum alleviates periodontitis. J Oral Microbiol 16:2388900.

113. Yang M, Dong P-T, Cen L, Shi W, He X, Li J. 2023. Targeting Fusobacterium nucleatum through chemical modifications of host-derived transfer RNA fragments. ISME J 17:880–890.

114. Vialetto E, Miele S, Goren MG, Yu J, Yu Y, Collias D, Beamud B, Osbelt L, Lourenço M, Strowig T, Brisse S, Barquist L, Qimron U, Bikard D, Beisel CL. 2024. Systematic interrogation of CRISPR antimicrobials in Klebsiella pneumoniae reveals nuclease-, guide- and strain-dependent features influencing antimicrobial activity. Nucleic Acids Res 52:6079–6091.

115. Doron L, Coppenhagen-Glazer S, Ibrahim Y, Eini A, Naor R, Rosen G, Bachrach G. 2014. Identification and characterization of fusolisin, the Fusobacterium nucleatum autotransporter serine protease. PloS One 9:e111329.

116. Vacca F, Sala C, Rappuoli R. 2022. Monoclonal Antibodies for Bacterial Pathogens: Mechanisms of Action and Engineering Approaches for Enhanced Effector Functions. Biomedicines 10:2126.

117. Richard JP, Melikov K, Vives E, Ramos C, Verbeure B, Gait MJ, Chernomordik LV, Lebleu B. 2003. Cell-penetrating Peptides: A REEVALUATION OF THE MECHANISM OF CELLULAR UPTAKE*. J Biol Chem 278:585–590.

118. Cockerill FR. 2012. Methods for dilution antimicrobial susceptibility tests for bacteria that grow aerobically: approved standard9th ed. Clinical and Laboratory Standards Institute, Wayne, Pa.

